# Found in translation: CCA-adding enzymes reveal bacterialization of Asgard archaea prior to eukaryogenesis

**DOI:** 10.64898/2026.05.29.728689

**Authors:** Ella Cassidy, Claudius Doktor, Heike Betat, Mario Mörl, Sonja J. Prohaska

**Author notes:** Authors contributed equally (Ella Cassidy, Claudius Doktor).

## Abstract

CCA-adding enzymes are essential for transfer RNA (tRNA) maturation and translation among all domains of life. A long-standing dogma holds that archaea encode exclusively class one CCA-adding enzymes (CCA1), whereas bacteria and eukaryotes encode class two enzymes (CCA2), creating a perceived strict evolutionary separation between archaeal and eukaryotic CCA-adding enzymes. Here, we show that this view is incorrect. We identify widespread CCA2 enzymes across multiple archaeal superphyla, including Asgard archaea, and demonstrate that these enzymes are functional enzymes and can replace their archaeal CCA1 counterparts. Within Heimdallarchaeia, the proposed closest archaeal relatives of eukaryotes, we identify distinct clades with different CCA-adding enzyme repertoires that trace back to independent bacterial sources. One Heimdallarchaeia lineage exhibits strongest similarity to Bacteroidota/Chlorobiota/Rhodothermota, a bacterial group previously implicated in early gene contributions to eukaryogenesis. CCA-adding enzymes from early-branching eukaryotes share this deep signature, indicating that predecessors of the eukaryotic enzymes had already been long-established and functionally adapted within the ancestral Asgardian gene pool. Together, these findings challenge a central paradigm of CCA-adding enzyme evolution and provide molecular-level support for the ancestral integration and subsequent transmission of bacterial functions prior to or during eukaryogenesis, aligning with current evolutionary frameworks such as the Heimdall nucleation–decentralized innovation–hierarchical import (HDH) model.

## Introduction

Identifying archaeal precursors of eukaryotic informational core systems is central to understanding eukaryote origins. Asgard archaea are widely regarded as the closest relatives of eukaryotes^1–5^. Their genomes appear to contain a mixture of archaeal core genes with substantial bacterial-derived content^3^. This mosaic architecture challenges the 40-year paradigm that the informational core of the cell, including transcription and translation, is exclusively archaeal in origin^4^. Instead, recent models like the Heimdall nucleation–decentralized innovation–hierarchical import (HDH) model^6^, suggest that a Heimdallarchaeia-like proto-eukaryotic host was a master collector of genetic material, stabilizing bacterial innovations prior to mitochondrial endosymbiosis. These observations support the idea that Asgard archaea acted as intermediates for the transmission of bacterial innovations into the proto-eukaryotic lineage. However, direct molecular evidence for how specific core cellular systems were transferred and stabilized during this transition remains elusive. The evolutionary origin of the eukaryotic CCA-adding enzyme represents one such unresolved puzzle.

Heimdallarchaeia are considered the archaeal group most closely related to eukaryotes, and serve as a key system for investigating evolutionary transitions^9^. Recent genome-scale analyses indicate that the assembly of the eukaryotic ancestor involved a sustained process of serial innovations that reshaped the Asgardian host^10^. Some studies suggest that this process was driven by a combination of acquisition of bacterial-derived functions, internal bioenergetic adaptations, or oxygen adaptations^11^. Ancient mobile genetic element (MGE) highways between certain Heimdallarchaeia families and bacterial clades (such as Bacteroidetes/Chlorobi) have been posited as a potential source of continuous importation of bacterial genes into Asgardian clades, predating eukaryogenesis^6^. Although the precise placement of eukaryotes within Heimdallarchaeia remains debated^12^, members of this clade consistently emerges as the closest archaeal relatives to eukaryotes^3,4,10^.

A potential marker for tracing these ancient transitions is the CCA-adding enzymes (also referred to as tRNA nucleotidyltransferase). It catalyzes the post-transcriptional addition of the CCA trinucleotide sequence to tRNA 3’-termini, a universal feature required for aminoacylation and translation^13^. Due to this essential function in protein biosynthesis, the CCA-adding enzyme is subject to strong purifying selection and retains deep evolutionary signals. It is discussed as an ancestral catalytic activity that exists in two structurally and mechanistically distinct classes^14–17^. Class 1 enzymes (CCA1) are found in archaea^18^, and class 2 enzymes (CCA2), in bacteria and eukaryotes (Fig. 1B)^19^. Within eukaryotes, CCA2s are grouped into: an alphaproteobacterial-type enzyme (aCCA) present in animals and acquired by horizontal gene transfer at the base of Holozoa, and an ancestral eukaryotic-type enzyme (eCCA) retained across all other eukaryotes (Fig. 1B)^8^, rendering eCCA the default form in eukaryotes. The origin of eCCA, however, remains unresolved. Since translation cannot proceed without a functional CCA-adding enzyme, eCCA must have been present at the base of the eukaryotic lineage. Its divergence from endosymbiotic bacterial enzymes and universal distribution across non-animal eukaryotes suggest an ancient origin predating the last eukaryotic common ancestor (LECA). Yet, no archaeal or bacterial progenitor has been identified so far.

**Figure 1.**
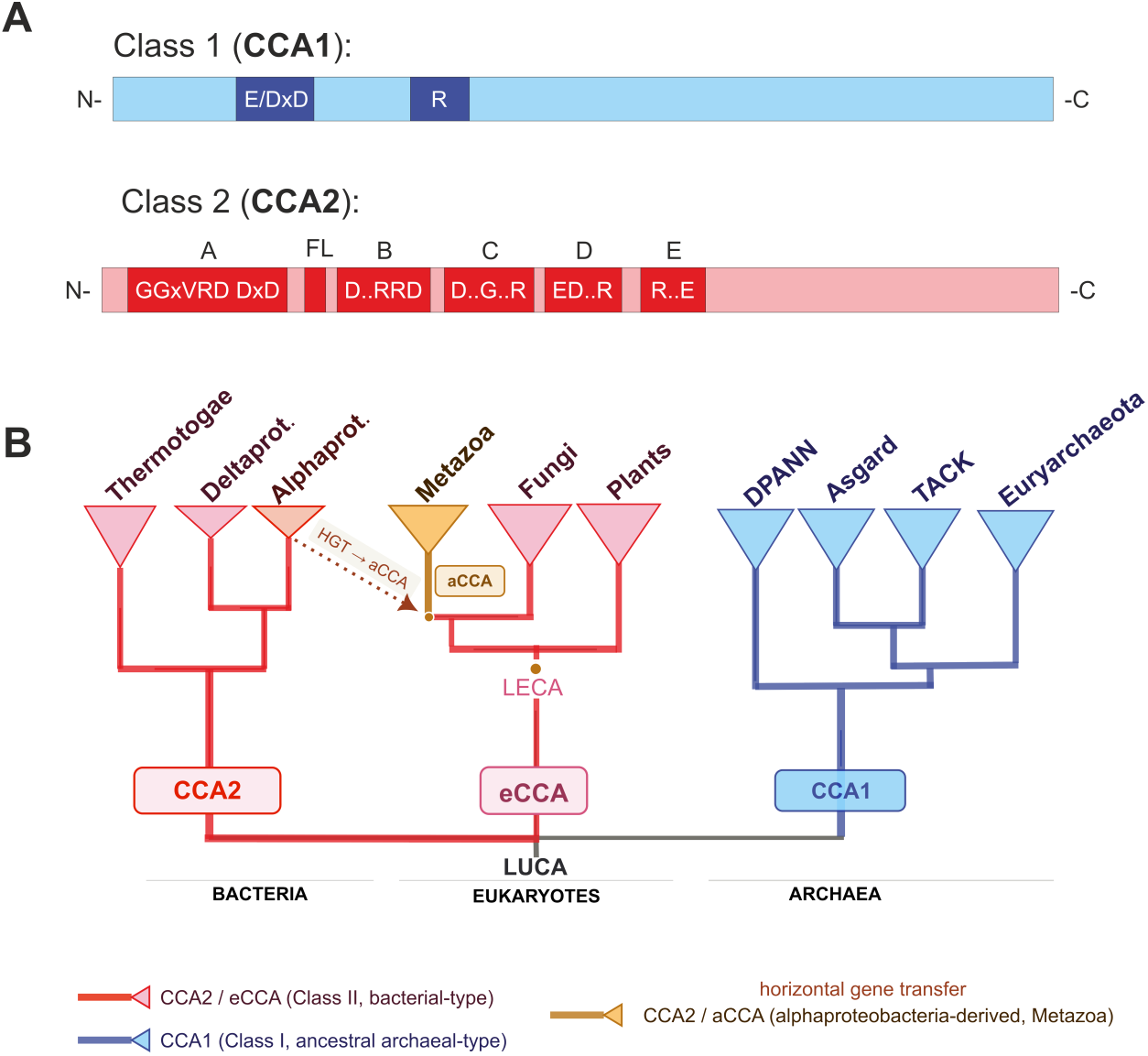
Structural architecture and evolutionary classes of CCA-adding enzymes. **A.** Schematic representation of the two major classes of CCA-adding enzymes and their conserved domain architectures. Both classes (class 1 and class 2) share the catalytically relevant sequence motif E/DxD or DxD motif (E, glutamate; D, aspartate; x, any amino acid)^7^. In class 1 (blue), a highly conserved arginine residue (R) is involved in CTP and ATP selection. Class 2 enzymes (red) are characterized by five conserved catalytic motifs (A - E) and a flexible loop element (FL) that is involved in the specificity switch from CTP to ATP. In motif D, residues EDxxR have templating function and specifically bind CTP and ATP by Watson-Crick-like hydrogen bonds. **B**. Traditionally, class 1 enzymes (CCA1) are restricted to Archaea, and class 2 enzymes to Bacteria and Eukarya. For CCA2, two subtypes are identified - an ancestral eukaryotic type (eCCA) which is found in plants, fungi and early branching eukaryotes, and an alphaproteobacterial-derived type (aCCA), which is found in animals^8^.

As a monomeric, essential catalyst, the CCA-adding enzyme retains deep evolutionary signals unavailable to multi-subunit systems. Here we exploit this property to demonstrate the presence of CCA2 in archaea and that bacterial-type enzymes were functionally integrated into the Asgardian translation core prior to eukaryogenesis and to verify functional activity of Asgardian CCA2 enzymes, providing an example of how a bacterial-derived informational component may have been stabilized prior to or during eukaryogenesis.

## Results

### Class 2 CCA-adding enzymes are widespread across archaeal lineages

Archaea were long thought to encode exclusively class 1 CCA-adding enzymes (CCA1). However, we identified class 2 CCA-adding enzymes (CCA2) across three of the four archaeal superphyla; Asgard, DPANN, and Euryarchaeota. Only TACK entirely lack CCA2 (Fig. 2). Since many of these genomes derive from metagenome-assembled genomes (MAGs), stringent quality controls were applied (≥85% completeness, <5% contamination) and duplicate genomes were removed prior to analysis. By applying these thresholds, we reduce the risk of chimeric assemblies that have been shown to overestimate the apparent eukaryotic affinity of certain Asgard lineages^2^, ensuring that the identified CCA2 sequences are not a result of contamination (full genome accessions, metadata, and CCA classifications for all retained archaeal genomes post-filtering are provided in Supplementary Table 1).

**Figure 2.**
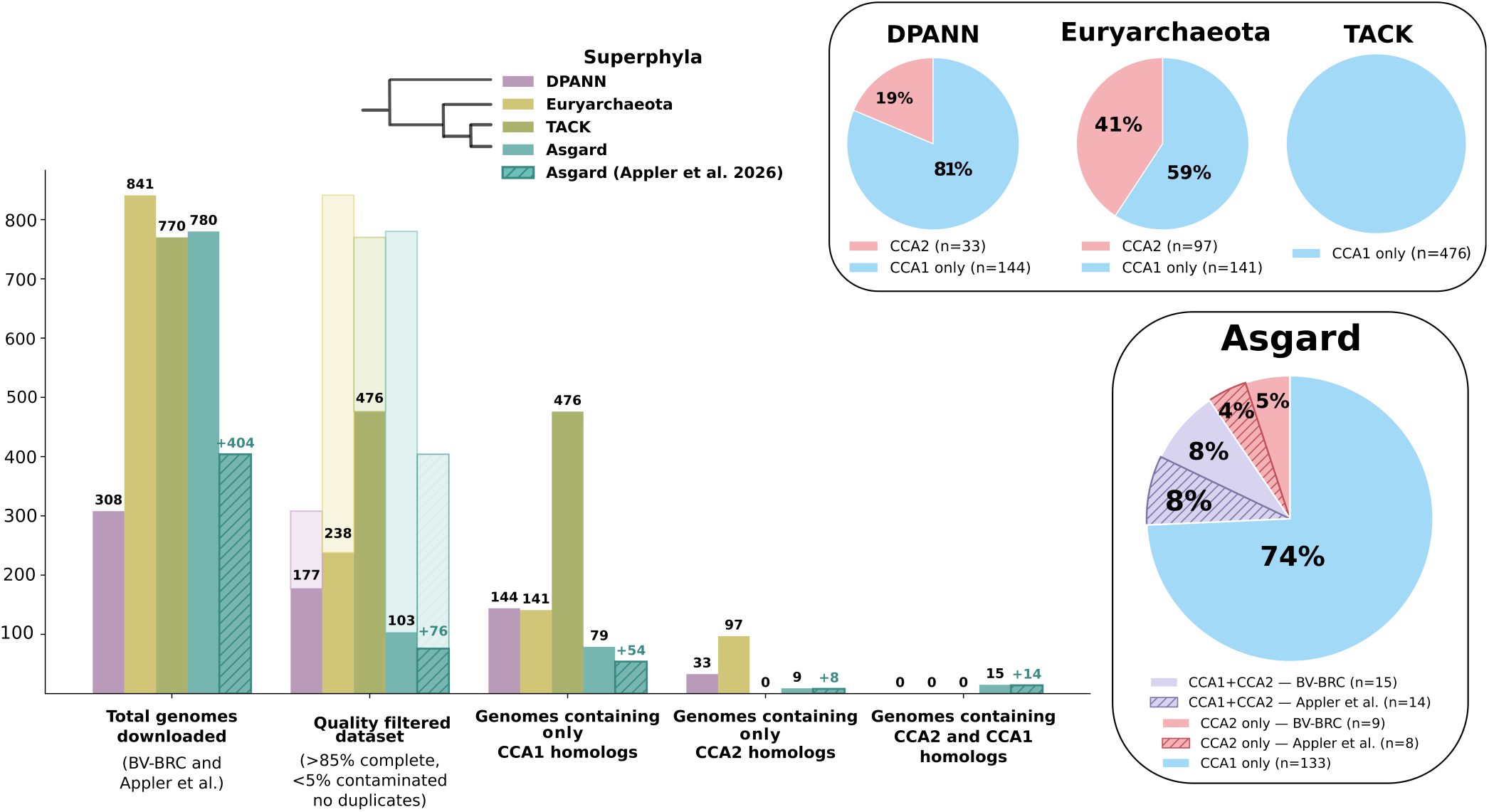
Overview of archaeal genome selection and CCA-adding enzyme classification. **A.** Schematic summary showing the total number of archaeal genomes analyzed, the number retained after quality filtering, and the distribution of class 1 (CCA1) and class 2 (CCA2) homologs across retained genomes (using the quality filtered dataset). **B.** Pie charts indicate the percentage of genomes encoding CCA2 relative to the quality filtered dataset. Colored sectors denote genomes containing CCA2 homologs (red), whereas blue sectors indicate genomes encoding CCA1 only, purple sectors denote the genomes percentage of genomes containing both CCA1 and CCA2.

To investigate whether archaeal CCA2 reflect coherent evolutionary histories or scattered acquisitions, we reconstructed species trees for Asgard, DPANN, and Euryarchaeota (using the genomes in the quality filtered dataset as seen in Fig. 2) using REvolutionH-tl^20^. REvolutionH-tl is a graph-theory-based phylogenomic method that infers orthogroups, event-labelled gene trees, and reconciled species trees directly from proteomes without relying on predefined marker gene sets. As TACK archaea only encode CCA1 and entirely lack CCA2, no species tree was constructed for this superphylum. In both DPANN and Euryarchaeota, CCA2-containing genomes clustered predominantly within monophyletic clades (Fig. 3), consistent with lineage-specific acquisitions rather than sporadic gene transfer events.

**Figure 3.**
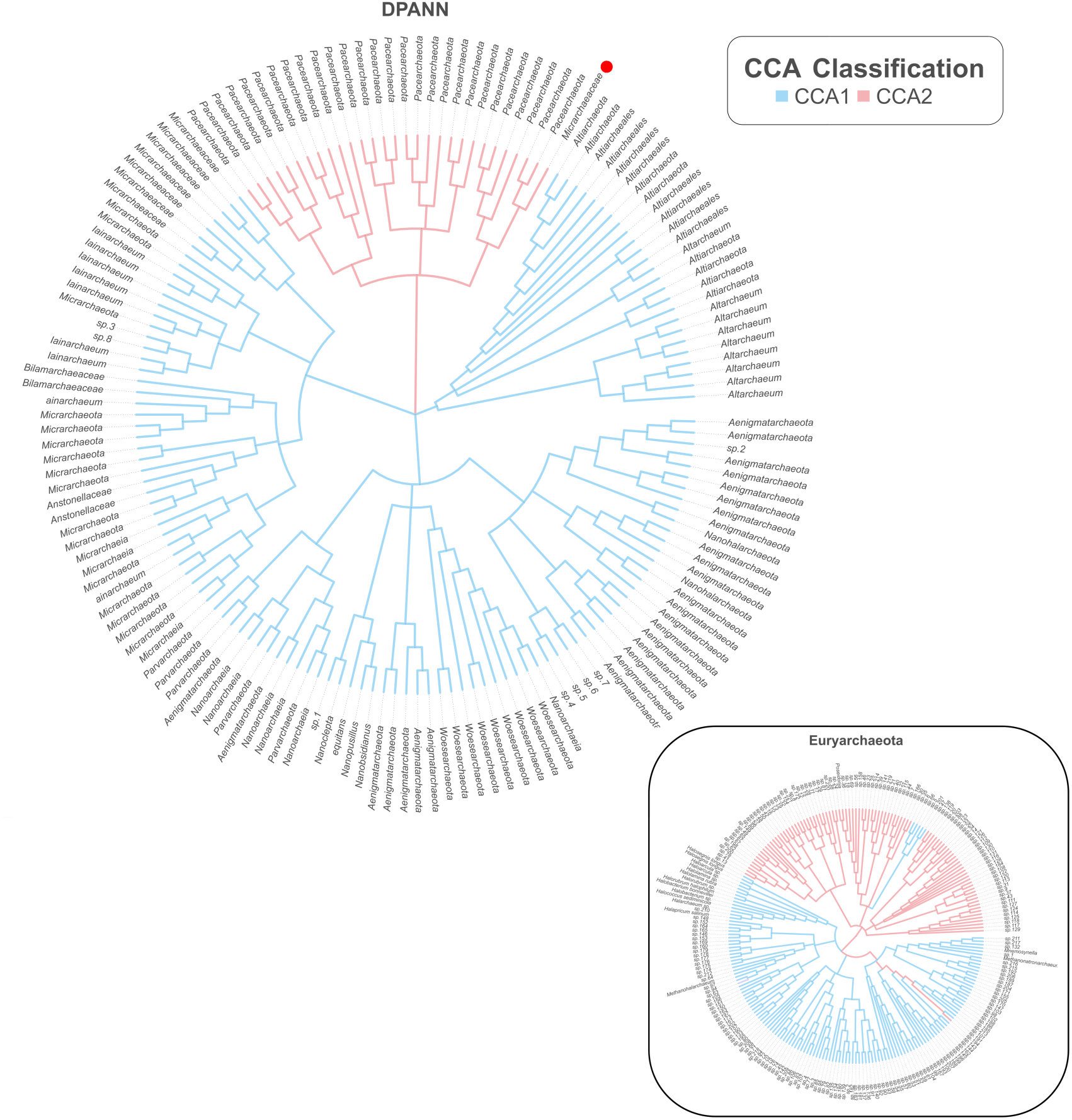
Species trees highlighting phylogenomic distribution of CCA-adding enzymes across DPANN and Euryarchaeota. Species trees reconstructed with REvolutionH-tl for DPANN (n=177) and Euryarchaeota (n=238) reveal the distribution of class 1 (CCA1, blue) and class 2 (CCA2, red) CCA-adding enzymes across major archaeal lineages. In DPANN, CCA2 is restricted to a monophyletic Pacearchaeota clade. In Euryarchaeota, CCA2 distribution is largely concordant with the species tree. See Supplementary Table 1 for details on genomes used in analysis with CCA class type. See Supplemental Figure x for an expanded Euryarchaeota species tree.

Within Euryarchaeota, CCA2 distribution is largely concordant with the species tree, forming monophyletic clades; however, five genomes (sp. 6, 47, 66, 201, and 5) encode only CCA1 form a monophyletic group yet nested within otherwise CCA2-containing clades, and one genome (sp. 194) encodes CCA2 despite falling within a CCA1 clade, suggesting secondary gene loss or horizontal transfer events in an otherwise stable distribution. In DPANN, CCA2 occurs exclusively in the monophyletic Pacearchaeota clade. A single Micrarchaeota genome (red dot, Fig. 3) is anomalously placed within this clade, while all other Micrarchaeota resolve in a separate part of the tree. Given that only high-quality MAGs were included (>85% complete, <5% complete according to CheckM2), and that REvolutionH-tl independently recovers the expected Micrarchaeota placement elsewhere, we interpret this genome as misbinned rather than a genuine instance of CCA2 acquisition in Micrarchaeota. Importantly, several DPANN and Euryarchaeota genomes encode CCA2 as their sole CCA-adding enzyme with no detectable CCA1 homologue. Since a functional CCA-adding enzyme is essential for tRNA maturation, this indicates that CCA2 is not a supplementary acquisition but has replaced the canonical class 1 enzyme as the active, functionally integrated system. Taken together, the concordance between CCA enzyme distribution and the species tree across both superphyla points to coherent vertical evolutionary histories, with isolated discordances reflecting rare rather than recurrent or recent horizontal acquisitions.

Among all archaeal groups, the Heimdallarchaeia clade in Asgards displayed the most diverse repertoire of CCA-adding enzymes, with genomes encoding CCA1 only, CCA2 only, or both enzyme classes simultaneously (Fig. 4). This distribution is unique within Asgard archaea, apart from Heimdallarchaeia, no other Asgard lineage encodes CCA2 enzymes. In connection with Heimdallarchaeia’s phylogenetic proximity to eukaryotes, this finding positions these Asgardian CCA2 enzymes as a focus for tracing the evolutionary origin of the ancestral eukaryotic CCA-adding enzyme (eCCA)^8^.

**Figure 4.**
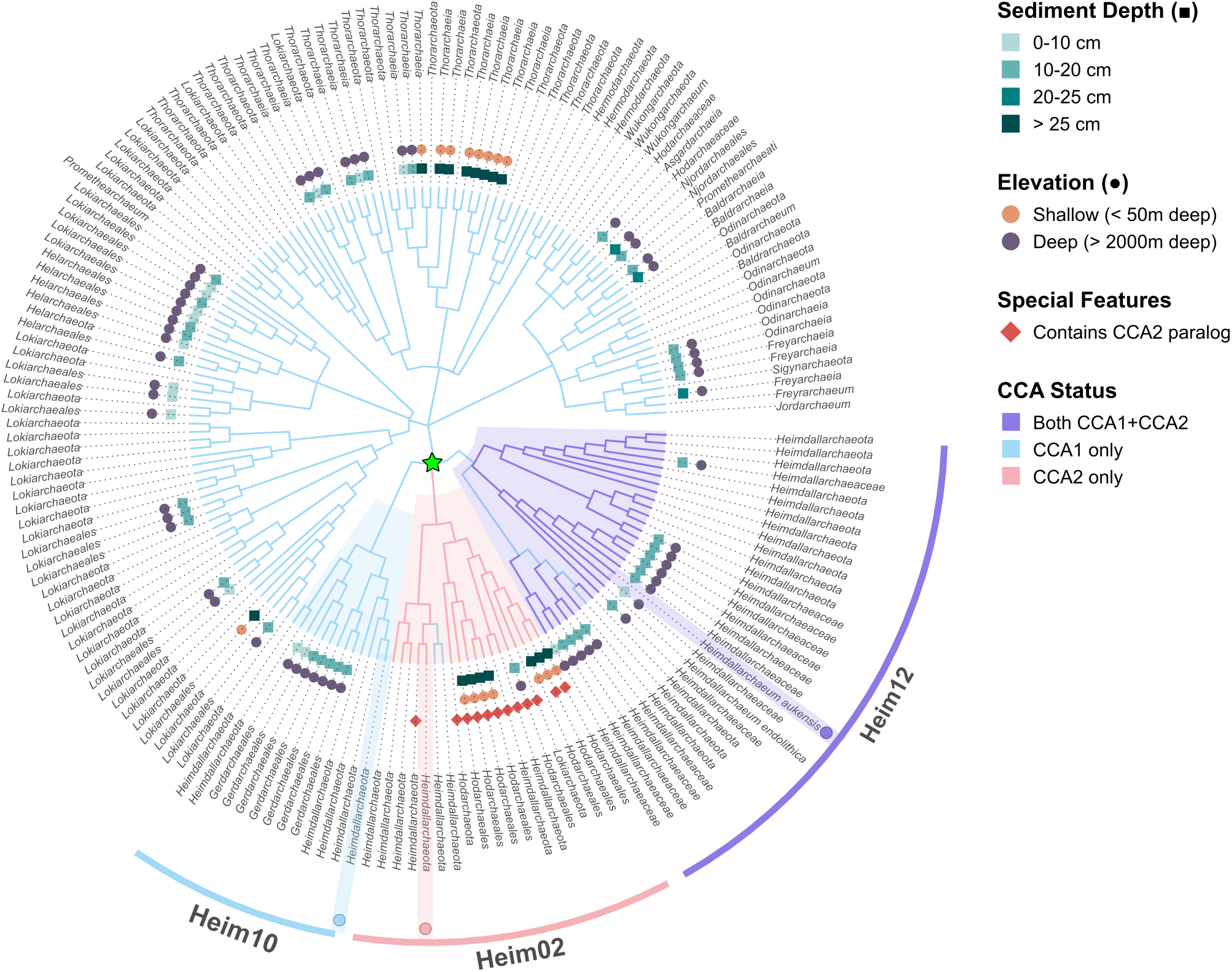
Evolutionary dynamics of CCA-adding enzymes in Heimdallarchaeia and other Asgard archaea. Species tree of the Asgard superphylum reconstructed with REvolutionH-tl^20^, showing the distribution of CCA-adding enzymes across major lineages. Branches are colored by CCA status. Three Heimdallarchaeia families are highlighted: Heim12 (Heimdallarchaeaceae, purple), encoding both CCA1 and CCA2; Heim02 (Hodarchaeales, red), encoding only CCA2; and Heim10 (Gerdarchaeales, blue), retaining only the ancestral CCA1. Metadata taken from Appler et al.^11^, including leaves are annotated with sediment depth (▪) and water column elevation (●) of the isolation site. Genomes within Heim02 that encode a CCA2 paralog are indicated by a red diamond (◆). The ancestral node of Heimdallarchaeia (encompassing Heim12, Heim02, and Heim10) is marked by a green star on the internal node.

Species tree topology and enzyme distribution revealed three distinct patterns across the major Heimdallarchaeia families: Gerdarchaeales encode only CCA1 enzymes (here labeled as Heim10), retaining the ancestral archaeal state; Heimdallarchaeaceae encode both CCA1 and CCA2 enzymes (here labeled as Heim12). Hodarchaeales carry only CCA2 enzymes (here labeled as Heim02), indicating that the ancestral CCA1 enzyme was replaced by a bacterial-derived CCA2 enzyme. Some Heim02 genomes additionally encode CCA2 paralogs, suggesting secondary duplication events following the initial replacement. The monophyly of Heimdallarchaeia was well-supported in REvolutionH-tl species trees, distinguishing this class from related but distinct Asgard lineages. Full species trees, including genome accession numbers, are provided in Supplementary Figure 2.

### Bacterial-type class 2 CCA-adding enzymes of Heimdallarchaeia are catalytically active and can replace archaeal class 1 CCA-adding enzymes

The phylogenetic distribution of class 1 and class 2 enzymes across the three domains of life was originally proposed in 1996^16^, at a time when sequence data was far more limited than today. With expanded genomic datasets available, CCA2 enzymes appear considerably more widespread in archaea than initially recognised, raising the question of whether these newly identified sequences represent functionally active enzymes subjected to selection and evolution, or merely conserved open reading frames. To address this, we selected CCA1 and CCA2 sequences from three Heimdallarchaeia species retrieved from the BV-BRC database: a *Gerdarchaeum* sp. representative of Heim10, placed within the Gerdarchaeales clade and encoding only CCA1; a *Hodarchaeum* sp. representative of Heim02, placed within the Hodarchaeales clade and encoding only CCA2; and *Heimdallarchaeum aukensis* representing Heim12, encoding both CCA1 and CCA2 (see Fig. 4 highlighted genomes selected, genome accessions provided in Supplementary Fig. 2). The coding sequences were inserted into plasmid pET-28a(+) and recombinantly expressed in *E. coli* BL21(DE3) Δ*cca*. The purified enzymes were tested for activity on a radioactively labeled yeast tRNA^Phe^ lacking the CCA-end (Fig. 5). While the enzymes from the Heim10 and Heim02 representatives showed the expected catalytic activity and added the CCA triplet, the class 1 enzyme from *H. aukensis* (Heim12) turned out to be completely inactive. A likely reason for this is that conserved regions accumulated inactivating mutations. As an example, the essential arginine residue involved in NTP selection in the catalytic core^21^, is replaced by asparagine, a strong indication that this enzyme has lost its CCA-adding activity. In addition, several insertions and deletions in comparison to canonical class 1 enzymes further support a functional inactivation of the protein (Supplementary Fig. 3). Inspection of CCA1 sequences across other Heim12 genomes reveals conservation of the key substitutions responsible for inactivity in *H. aukensis*, suggesting that enzymatic inactivity of the class 1 enzyme is a general feature of Heim12 rather than a lineage-specific anomaly. The corresponding class 2 enzyme, however, was fully active on the tRNA substrate and incorporated a complete CCA-end. Taken together, these results indicate that a class 2 enzyme can indeed replace the activity of the ancestral class 1 enzyme and represent a bona fide and functional CCA-adding enzyme.

**Figure 5.**
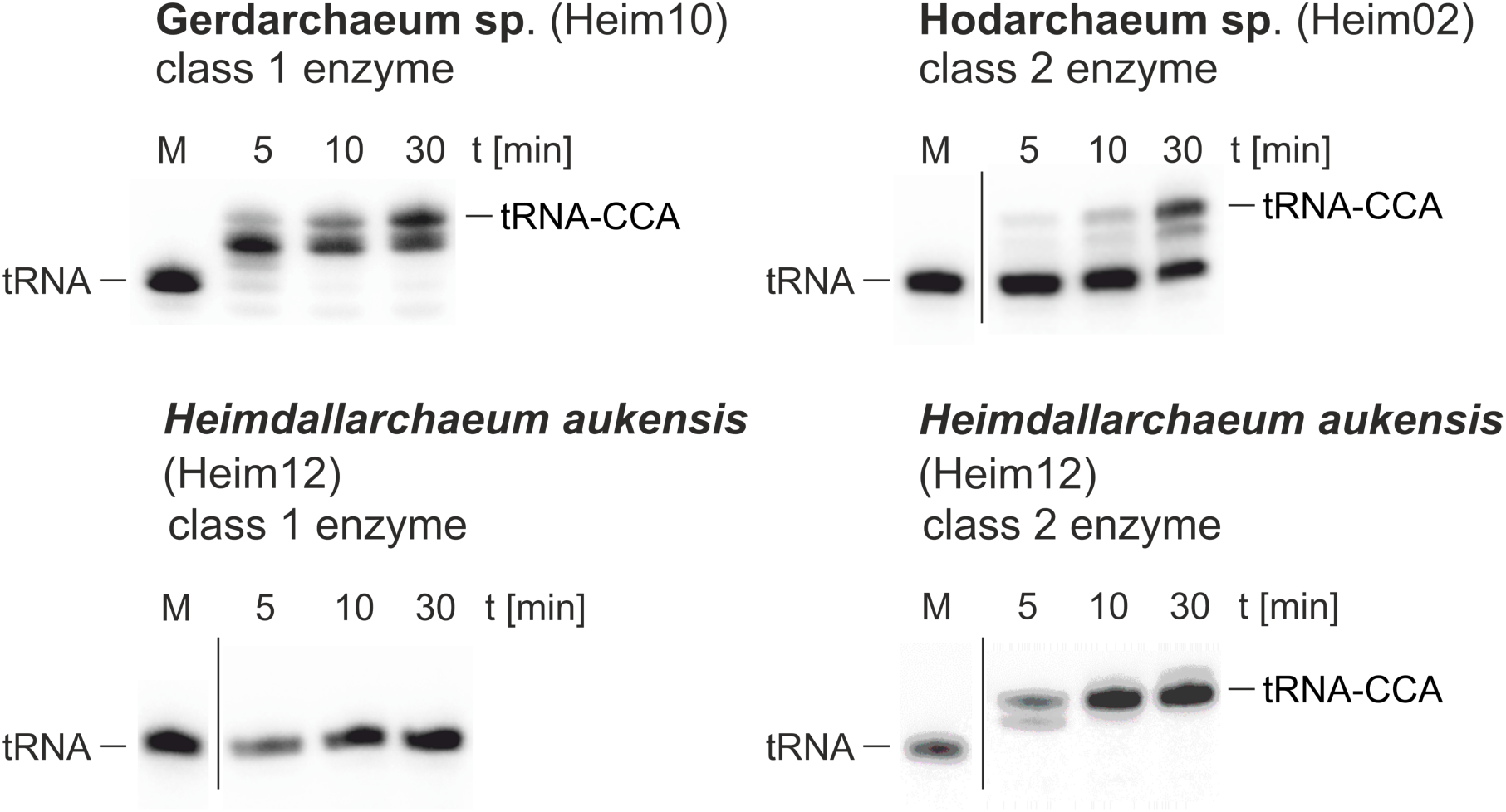
Activity tests of recombinant CCA1 and CCA2 enzymes from selected Heimdallarchaeia species. CCA-addition on tRNA substrate. In a time series experiment, the recombinant class 1 enzyme from Gerdarchaeuum sp. shows a complete CCA incorporation, although the addition of the terminal A position was less efficient (upper left panel). This is probably due to non-optimal *in vitro* incubation conditions for this protein. Upper right panel: the class 2 enzyme from Hodarchaeum sp. exhibits full CCA-addition. In contrast, the class 1 enzyme from Heim12 Heimdallarchaeum aukensis is completely inactive (lower left panel), while the corresponding class 2 enzyme obviously took over the activity and incorporates a complete CCA triplet (lower right panel).

### Phylogenetic Analysis of CCA2 Enzyme Evolution in Bacteria

We first considered whether CCA2 could represent an ancestral archaeal feature or instead is a later bacterial-derived acquisition. Although CCA2 homologues occur in DPANN, Asgards and Euryarchaeota, lineage-restricted distributions and occasional discordance with the species tree are more consistent with horizontal gene transfer. In contrast, TACK archaea entirely lack CCA2 sequences across all retained genomes, which makes a single archaeal ancestral origin unlikely. Explaining the observed distribution by vertical inheritance would require multiple independent losses across one of the most diverse archaeal superphyla, a scenario less parsimonious than lineage-specific bacterial-to-archaeal transfers. Therefore, we interpret all archaeal CCA2 sequences as bacterial-derived acquisitions.

To identify the bacterial donors of Heimdallarchaeia CCA2 enzymes and to place these transfer events into an evolutionary context, we characterized the diversity and evolutionary structure of bacterial CCA2 sequences across a representative bacterial dataset (see Supplement Table 2 for details about bacterial groups, including genome identifiers). This analysis revealed two major clusters that reflect different evolutionary histories (Fig. 6). These clusters are referred to here as early- and late-diverging, defined not by their position in the bacterial species tree but by the GOE-relative metabolic identity of their constituent lineages - that is, whether their core metabolic lifestyle predates or postdates atmospheric oxygenation. The early-diverging cluster includes lineages with pre-GOE origins or anaerobic/thermophilic metabolic identities. This cluster spans deep-branching thermophiles (Aquificota, Chloroflexota, Thermotogota, Deinococcota, and Caldisericota) and anaerobic lineages (Thermodesulfobacteriota and Elusimicrobiota). Also resolving within this cluster are the Bacteroidota, Chlorobiota and Rhodothermota phyla, whose CCA2 sequences retain ancestral signatures consistent with an ancient stem-group origin^22^. A key feature of this cluster is the frequent occurrence of subfunctionalized variants: rather than encoding a single CCA-adding enzyme, many of these lineages encode separate CC-adding and A-adding enzymes that collaborate in synthesizing a complete CCA-end. Identification of these variants is possible due to structural hallmarks: bacterial CC-adding enzymes are characterised by a deletion of the flexible loop element, which inhibits the specificity switch from CTP to ATP^17^, along with A-adding enzymes containing a downstream motif sRxxxExxxhh^19^.

**Figure 6.**
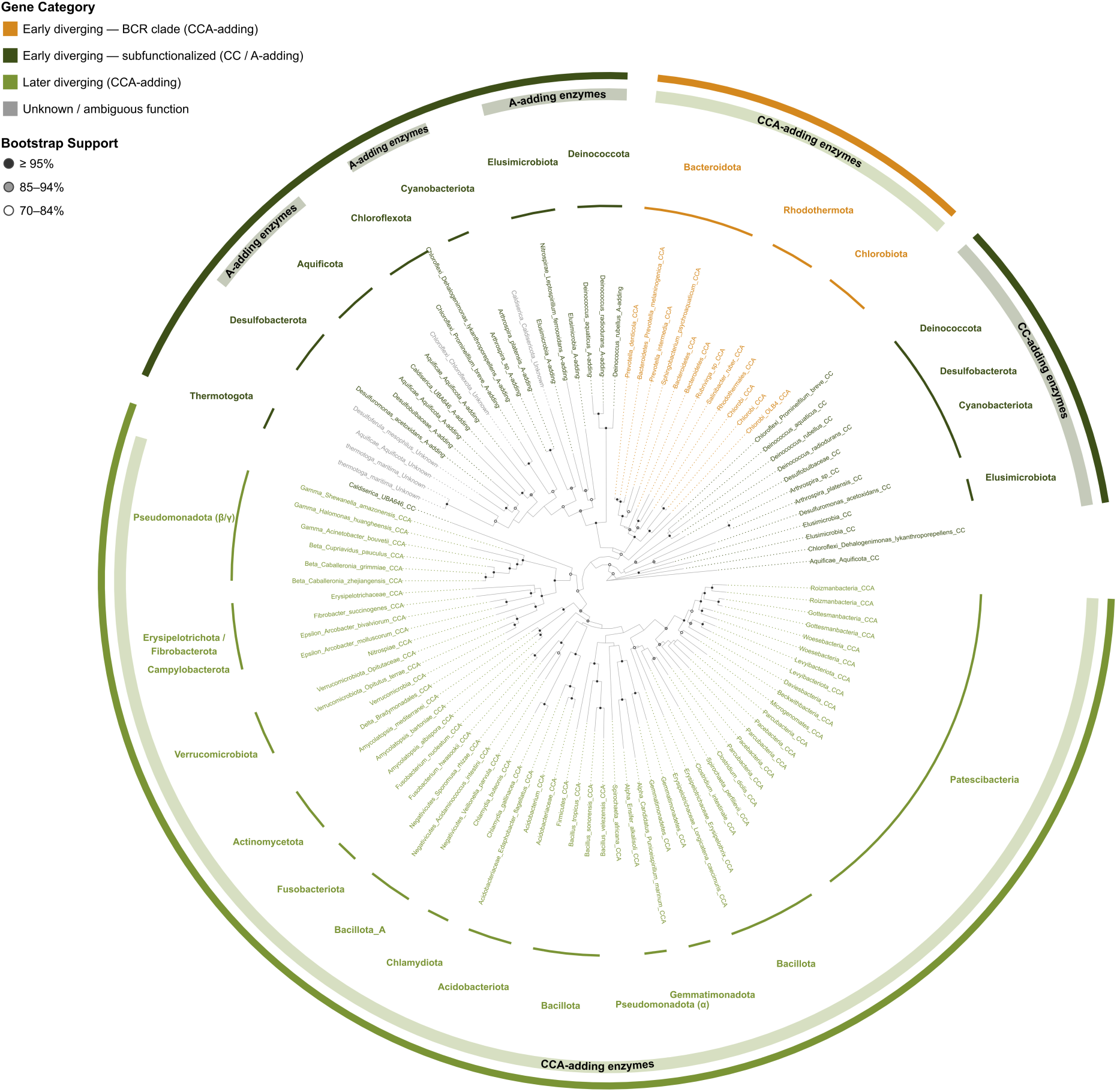
Phylogenetic diversification and subfunctionalization of class 2 CCA-adding enzymes in Bacteria. Maximum- likelihood gene tree of bacterial class 2 tRNA nucleotidyltransferases inferred using IQ-TREE 3 with ModelFinder, under the Q.PFAM+I+R6 substitution model selected according to the Bayesian Information Criterion (BIC). Node support values represent ultrafast bootstrap (UFBoot) proportions from 1000 replicates. The tree is rooted using Aquificota as outgroup. Two major clusters are identified based on enzyme architecture and ecological identity relative to the Great Oxidation Event (GOE). The early-diverging cluster (dark green) encompasses lineages with pre-GOE metabolic identities, including thermophilic and anaerobic taxa, and is characterised by frequent subfunctionalization. Many lineages encode separate CC-adding and A-adding enzymes rather than a single bifunctional CCA-adding enzyme. The Bacteroidota/Chlorobiota/Rhodothermota superphylum (yellow) resolves within this cluster, retaining canonical CCA-adding enzymes. The later-diverging cluster (light green) encompasses lineages associated with the post-GOE aerobic radiation and exclusively encodes canonical full-length CCA-adding enzymes. Bacterial group nomenclature follows the GTDB taxonomy^35^.

By screening for these structural hallmarks, subfunctionalized enzyme pairs can be identified and distinguished from canonical single-enzyme CCA-adding activity (see Methods). This subfunctionalization has been identified and biochemically validated in bacterial groups including Deinococcota, Cyanobacteriota, and Aquificota, where CC- and A-adding enzymes collaborate to construct the complete CCA terminus^23–26^. Our analysis indicates that this enzymatic collaboration is more widespread in these early-diverging lineages than previously recognised. Notably, all Thermodesulfobacteriota sequences in our dataset (spanning both thermophilic lineages and anaerobic sulfate- and sulfur-reducing members previously classified within Deltaproteobacteria, including Desulfobulbaceae and Desulfuromonas^27^), are subfunctionalized and placed within the early-diverging cluster. Elusimicrobiota, obligate anaerobic endosymbionts with restricted metabolic capacity, likewise contribute both CC- and A-adding variants to this cluster. Yet, in certain lineages within this cluster, including Bacteroidota, Chlorobiota, and Rhodothermota, canonical CCA-adding enzymes are retained. Despite the Chlorobiota crown group radiating relatively recently at ∼1.88 Gya^28^, CCA2 sequences from all three phyla retain ancestral signatures shared with the thermophilic and deep-branching subfunctionalized lineages in this cluster, consistent with a stem-group origin predating their divergence.

The second later-diverging cluster groups lineages with post-GOE metabolic identities, predominantly aerobic or facultatively aerobic organisms whose major ecological diversification was enabled by atmospheric oxygenation. This group includes Pseudomonadota, Bacillota, Spirochaetota, Acidobacteriota, Verrucomicrobiota, Fibrobacterota, Actinomycetota, and Patescibacteria (formerly CPR)^29–32^. In contrast to the early-diverging cluster, this bacterial cluster exclusively encodes canonical CCA-adding enzymes, including the aCCA-type lineages found in Alphaproteobacteria (Pseudomonadota α). The contrasting placement of Thermodesulfobacteriota and Bdellovibrionota, two lineages historically classified within Deltaproteobacteria, is particularly informative in this context: while anaerobic sulfate- and sulfur-reducing Thermodesulfobacteriota are placed within the early-diverging cluster, aerobic predatory Bdellovibrionota resolve within the later-diverging cluster, reflecting their contrasting metabolic identities. Although the Patescibacteria divergence ages are subject to debate^33^, Patescibacteria sequences cluster with these later-diverging bacterial groups despite the possible ancient origin of this radiation.

Taken together, these two bacterial CCA2 clusters, grounded in the subfunctionalization pattern of CC- and A-adding enzymes and corroborated by geological constraints, provide a useful temporal reference for interpreting the antiquity of bacterial-to-archaeal CCA2 transfer events. Although this two-cluster structure does not recapitulate the canonical Terrabacteria/Gracilicutes split, it remains a useful evolutionary tracer. Rather than tracking vertical organismal descent, CCA2 acts as a functional and temporal tracer: structural innovations within this family reflect host metabolic ecology relative to the GOE, providing a reliable reference for the timing of ancient transfers^22,34^.

Using this framework, we interpreted the archaeal CCA2 gene tree in terms of relative evolutionary timing regarding the GOE to better understand the ancient horizontal gene transfers into archaeal lineages (Fig. 6). The root placement is consistent across multiple methods. Midpoint rooting places Aquificota and Arthrospira CC-adding enzymes at the base, and Minimal Ancestor Deviation (MAD) rooting independently identifies a CC-adding enzyme from Caldisericota, converging on subfunctionalized CC-adding enzymes from thermophilic lineages regardless of approach.

### Divergent Bacterial Origins of Heimdallarchaeia CCA2 enzymes

Pairwise comparisons of Heimdallarchaeia CCA2 enzyme sequences against the full bacterial dataset revealed a clear separation between Hodarchaeales (Heim02) and Heimdallarchaeaceae (Heim12) (Fig. 7). Hodarchaeales homologs showed strongest similarity to later-diverging bacterial lineages (Patescibacteria, Gemmatimonadetes, Bacillus, Fusobacterium, and Clostridium), all members of the post-GOE radiation. This affiliation with the later-diverging cluster is consistent with a horizontal transfer from a post-GOE bacterial donor. Notably, several Hodarchaeales genomes encode two CCA2 copies, denoted as A and B paralog. The lack of archaeal CCA1 in Hodarchaeota genomes shows a complete replacement of their original archaeal system and functional replacement. The species tree reveals an internal split within Hodarchaeales that follows paralog presence or absence (Fig. 4), suggesting either an intra-lineage duplication after the initial acquisition, or two independent transfers.

**Figure 7.**
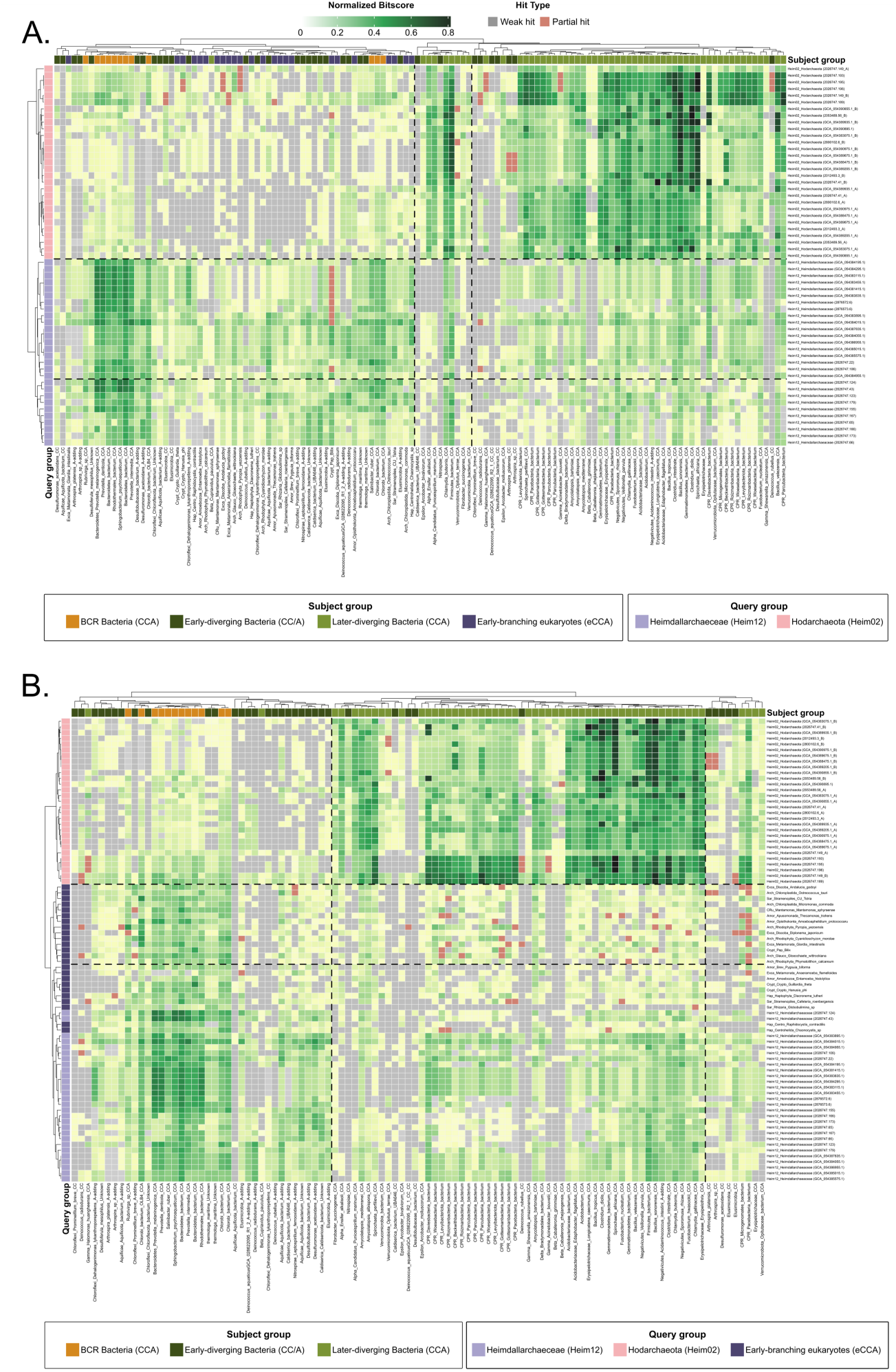
Divergent evolutionary origins of Heimdallarchaeia CCA2 and shared ancestral bacterial signature with early-branching eukaryotes. Pairwise BLASTp bit scores. Color intensity corresponds to BLAST bit score normalized by alignment length (see Methods), where grey cells are weak hits with good coverage (weak hit), and red cells are good hits with short alignment lengths (partial hit). In panel A., Heimdallarchaeaceae (Heim12) sequences (purple, y-axis) show highest similarity to Bacteroidota/Chlorobiota/Rhodothermota (orange, x-axis), an early-diverging bacterial superphylum with pre-GOE ancestral origins, whereas Hodarchaeales (Heim02) sequences (pink, y-axis) show highest similarity to later-diverging bacterial lineages including patescibacteria, Firmicutes, and Proteobacteria (light green, x-axis). In panel B., when early-branching (EB) eukaryotic eCCA sequences are included in the query set alongside Heimdallarchaeia sequences, EB eukaryotes and Heimdallarchaeaceae (Heim12) (both purple, y-axis) converge on the same Bacteroidota/Chlorobiota/Rhodothermota (BCR) affiliation, while Hodarchaeales (Heim02) retains its later-diverging bacterial signature.

Heimdallarchaeaceae (Heim12) sequences showed a fundamentally different pattern, aligning most closely with Bacteroidota, Chlorobiota, and Rhodothermota (BCR) members of the early-diverging bacterial cluster (Fig. 7a and 7b). While the crown group of Chlorobi radiated after the GOE (∼1.88 Gya)^28^, its last common ancestor with Bacteroidota and Rhodothermota predates other bacterial lineages. The similarity of their CCA2 sequences suggests that they still share signatures of ancestral origin. The affiliation of Heim12 with this ancient bacterial superphylum, along with a more eroded signal strength, places the Heim12 acquisition deeply back in time, consistent with a pre-GOE or early-GOE transfer from a stem-group BCR ancestor. Together these patterns suggest that Hodarchaeales (Heim02) and Heimdallarchaeaceae (Heim12) acquired CCA2 sequence independently from distinct bacterial donors at different points in evolutionary time.

To determine whether either Heimdallarchaeia lineage shares its bacterial signature with eukaryotic eCCA enzymes, we expanded the BLASTp query set and included early-branching ancestral eukaryotic eCCA sequences alongside the Heimdallarchaeia sequences (Fig. 7b). The eCCA sequences showed highest similarity to Bacteroidota, Chlorobiota and Rhodothermota the same early-diverging bacterial lineages most similar to Heimdallarchaeaceae (Heim12), but not to the later-diverging lineages linked to Hodarchaeales (Heim02). This convergence is not driven by direct similarity between Heim12 CCA2 sequences and eukaryotic eCCA sequences, which is weak as expected after over more than two billion years of independent evolution of the CCA2 gene in archaeal and eukaryotic lineages. Rather, both Heim12 and early-branching eukaryotes independently point to the same bacterial donor group, with the bacterial sequences serving as the common reference that has not undergone the same degree of lineage-specific changes. This pattern is consistent with previous genome-scale studies suggesting that Bacteroidota and Chlorobiota contributed bacterial genes to Asgard archaeal ancestors of eukaryotes, and these acquisitions were subsequently stabilised in archaeal genomes prior to eukaryogenesis^6^. The homology signal strength (normalized bitscore) between BCR-clade CCA2 sequences and eukaryotic CCA2 sequences is comparable to that observed between the BCR-clade and Heim12, suggesting that both lineages have diverged from the bacterial donor over a similar evolutionary timespan. This shared, eroded bacterial signal in both Heimdallarchaeaceae and early-branching eukaryotes are consistent with a model in which a bacterial-type CCA2 enzyme was integrated into an ancestral Heimdallarchaeia framework prior to the divergence of the eukaryotic lineage, rather than representing a late, independent acquisition.

This evolutionary trajectory mirrors the deep mosaicism documented in the prokaryotic aminoacyl-tRNA synthetase (aaRS) repertoire, where ancient non-orthologous gene displacements resulted in the replacement of native archaeal enzymes with bacterial counterparts^36^. While these classic aaRS replacements represent deep prokaryotic inter-domain transfers, our findings demonstrate that a similar capacity for informational core overhauls occurred within the specific Asgard lineages ancestral to eukaryotes. Ultimately, the CCA2 enzyme provides a rare, highly resolved opportunity to observe this fundamental pre-mitochondrial ‘bacterialization’ of the host informational machinery within a single-copy essential gene. The feasibility of this ancient transfer is supported by evidence of an ancient and scalable viral highway between Asgard archaea and the BCR phylum. The presence of integrated viral genomes in Heimdallarchaeia (e.g., *HeimV1*) containing integrase genes closely related to Bacteroidetes-hosted bacteriophages identifies viral-mediated HGT as a primary, deep-time mechanism for hierarchical import^6^. While these mobile genetic elements (MGEs) provide the physical mechanism, the erosion of flanking signatures (such as transposases or microsynteny) at the Heim12 CCA2 locus points toward an ancient acquisition.

Reliable deep-time phylogenetic inference requires mitigating homoplasy and Long Branch Attraction (LBA) driven by fast-evolving, compositionally biased sites. Full-length CCA-adding enzymes possess a universally conserved catalytic core and a highly variable N- and C-termini (Fig. 1a). Evaluation of our full-length alignment (987 sites, 164 sequences) revealed that terminal regions are largely gap-filled, with 95.5% of terminal sites falling into the fastest-evolving categories (Supplementary Fig. 4, see Methods), suggesting these regions reflect alignment artefact rather than genuine phylogenetic signal. These gaps were visualized by calculating the proportion of sequences carrying a gap character at that position.

Consequently, 69.4% of sequences failed compositional homogeneity tests. While progressive gap-threshold filtering (KPIC-Gappy) yielded minimal improvement, maintaining 60.6% failure (Supplementary Fig. 4, see Supplementary Table 4), excising the variable termini substantially reduced compositional heterogeneity (34.4% failure) and removed the primary drivers of LBA risk. This structural truncation is biochemically justified, as these terminal regions are dispensable for catalytic activity^21^. To maximize phylogenetic signal and prevent artifactual topologies, the following analyses were restricted to the conserved catalytic core.

### A Deep Evolutionary Bridge to Eukarya

To reconstruct a reliable evolutionary history of the CCA-adding enzyme, we performed maximum-likelihood phylogenetic inference. Because deep phylogenies are notoriously susceptible to compositional bias and alignment noise, we applied stringent site-filtering (see Methods). This optimization mitigated compositional heterogeneity, reducing the proportion of sequences failing the chi-square test to 34.4% as reported by iqtree3^37^, and yielded an alignment of 205 amino acid sites. First, an initial maximum-likelihood tree was inferred using the Q.PFAM+I+R6 substitution model, selected via ModelFinder^38^, to account for conserved domain evolution. This initial topology was then utilized as a guide tree to estimate site-specific frequency profiles for a Posterior Mean Site Frequency (PMSF) analysis^39^, allowing us to model site-specific compositional heterogeneity. Both maximum-likelihood (ML) and Bayesian frameworks consistently recovered a robust, monophyletic clade uniting early-branching eukaryotes, Heimdallarchaeaceae (Heim12), and the bacterial Bacteroidota/Chlorobiota/Rhodothermota group (BCR); hereafter referred to as the EHB clade (Fig. 8; Supplementary Fig. 5). The monophyly of this broad group is strongly supported in the ML framework (UFBoot = 99%, SH-aLRT = 93%), and is independently corroborated by the Bayesian topology, which recovered the identical EHB clade despite non-convergence between independent runs (see Methods). This methodological congruence confirms that the EHB grouping is a genuine evolutionary signal rather than a reconstruction artefact such as long-branch attraction and is consistent with the convergent signal identified in our pairwise BLASTp analyses (Fig. 7b). To formally evaluate support for this arrangement, we tested all three possible topologies for the BCR, Heim12, and eukaryote clades using the approximately unbiased (AU) test. None of the three alternative arrangements could be significantly rejected (p-AU = 0.699, 0.426, and 0.179, respectively); the phylogenetic signal available in the 205-site catalytic core alignment is insufficient to statistically discriminate between them. Nevertheless, the topology placing BCR as sister to a (Heim12 + eukaryote) clade recovered the highest log-likelihood of the three (ΔlnL = 0), consistent with the PMSF bootstrap support.

**Figure 8.**
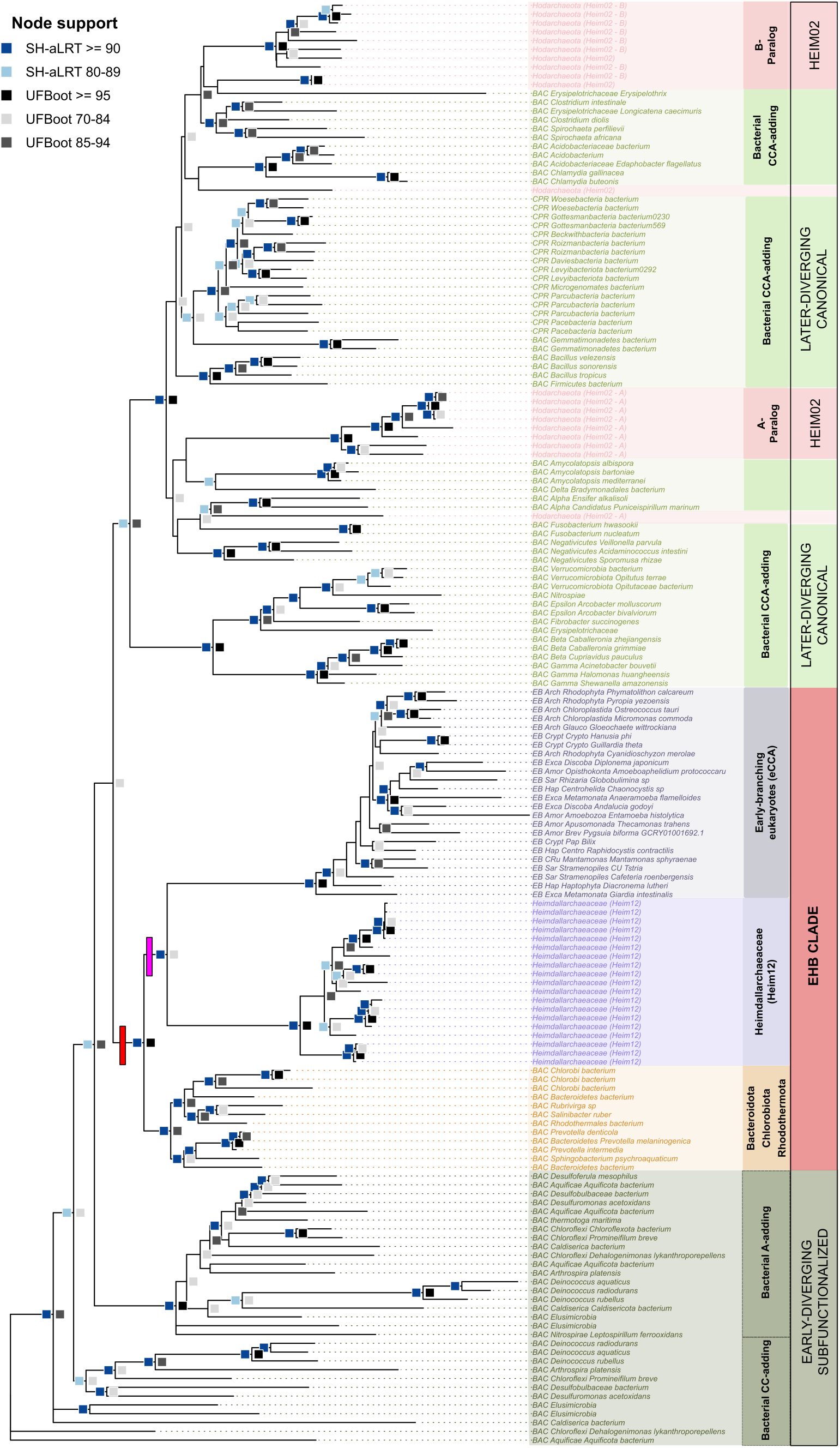
The eukaryotic, Heimdallarchaeaceae, and bacterial (EHB) CCA2 clade is supported across phylogenetic frameworks. Support values at nodes are indicated by square shading (see legend for UFBoot and SH-aLRT thresholds). The monophyly of the EHB clade is strongly supported in the ML framework (UFBoot > 99%; SH-aLRT > 93%). The red bar denotes the common ancestor of the EHB clade, whereas the pink bar denotes the inferred horizontal transfer event giving rise to the shared Heim12–eukaryote CCA2 lineage. Sequence annotations on the right indicate functional and taxonomic groupings: subfunctionalized CC- and A-adding enzymes are shown in dark green; canonical non-subfunctionalized CCA-adding enzymes in light green; early-branching eukaryotes (eCCA) in dark purple; Heimdallarchaeaceae (Heim12) in lavender, collectively forming the EHB clade with Bacteroidota/Chlorobiota/Rhodothermota (red); and Hodarchaeales (Heim02) in light pink, with B- and A-paralog pairs annotated separately. The Bayesian topology, presented as a supplementary figure (Supplementary Fig. 5), independently recovered the identical EHB clade (see Methods).

The tree exhibits a distinct branch-length hierarchy (eukaryotic branches are longest, followed by Heim12, with BCR showing the shortest stems, represented by red bar in Fig. 8), where the short stems of the bacterial BCR group represent the more ancestral bacterial state, and the elongated eukaryotic and Heimdallarchaeaceae branches reflect accelerated sequence divergence. Rather than an independent or recent lateral acquisition, this topology provides powerful molecular evidence that the bacterialization of the eukaryotic translation core was initiated via a pre-mitochondrial capture event in an Asgard ancestor over 2 billion years ago. The shared evolutionary trajectory of the CCA2 sequence in both Heim12 and early-branching eukaryotes supports the frameworks of the Serial Innovations and Heimdall nucleation– decentralized innovation–hierarchical import (HDH) models ^6,10^. The pink bar (Fig. 8) denotes the last common ancestor of Heimdallarchaeaceae and eukaryotes in this gene tree, representing the point at which the BCR-affiliated CCA2 enzyme, already established in the Heim12 lineage, was transferred into the proto-eukaryotic lineage, consistent with an archaeal rather than direct bacterial origin of the eukaryotic CCA-adding enzyme.

This gene tree shows a deep evolutionary split within the Asgard archaea. While the Heimdallarchaeaceae (Heim12) group is located firmly within the ancestral EHB clade, the Hodarchaeales (Heim02) cluster entirely outside this radiation, nesting instead within later-diverging bacteria (Fig. 8). This topological separation indicates that Hodarchaeales and Heimdallarchaeaceae acquired their class 2 CCA enzymes through separate evolutionary trajectories. This distinction is important for the current debate on eukaryotic origins: although species-tree analyses frequently place Hodarchaeales as the closest known archaeal relatives of eukaryotes, their class 2 CCA enzyme represents a late, post-GOE acquisition. In contrast, the Heimdallarchaeaceae preserve the ancient BCR-like profile that mirrors the eukaryotic state. Rather than implying a direct linear ancestry, this distribution aligns with the HDH model, where this enzyme represents a decentralized innovation from the ancient BCR radiation that was permanently stabilized within an ancestral Heimdallarchaeia framework to replace its native class 1 counterpart.

The antiquity of the Heim12 transfer is further corroborated by genomic and ecological context. The absence of localized microsynteny or mobile genetic element signatures flanking the Heim12 class 2 locus points to an ancient, integration event. Ecologically, modern Heimdallarchaeaceae and Bacteroidota persistently co-occur in bathypelagic marine sediments^40^, offering plausible environmental proximity that facilitated this exchange. Rather than a passive accumulation of foreign genes, the Heim12 ancestor underwent a fundamental reconfiguration of its translational apparatus. By swapping its native class 1 enzyme for a biochemically distinct class 2 variant, the lineage potentially optimized protein synthesis efficiency to complement the expanded bioenergetic capacities provided by its specialized proton pumps^11^.

## Discussion

Our findings provide a new perspective on the evolution of tRNA nucleotidyltransferases and the assembly of the proto-eukaryotic translation machinery. By identifying phylogenetically widespread CCA2 enzymes across three superphyla of archaea, and demonstrating biochemical functionality in Heimdallarchaeia, we show that bacterial-derived translation machinery can be stably integrated into archaeal lineages ancestral to eukaryotes, extending evidence for bacterialization into a component of the informational core previously considered to be of archaeal descent.

The results suggest that the ancestral eukaryotic enzyme (eCCA) originated as a bacterial-derived innovation in the Bacteroidota/Chlorobiota/Rhodothermota (BCR) stem lineage before being integrated and stabilized within an ancestral Heimdallarchaeaceae (Heim12) lineage. Recent global surveys have concluded that the translational machinery of eukaryotes is almost purely of Asgard archaeal descent^3^. The bacterial-derived CCA-adding enzyme represents a rare but functionally critical outlier to this pattern, suggesting that even the core protein synthesis machinery was subject to targeted bacterial innovation prior to eukaryogenesis. It is also consistent with the Heimdall nucleation (HDH) framework, which proposes that the Asgardian host accumulated multiple bacterial-derived innovations before the emergence of eukaryotes^6^. The HDH model initially addressed the decentralized assimilation of metabolic functions during early eukaryogenesis. This framework has since expanded: Feng et al.^10^ demonstrated that core eukaryotic replisome components, long viewed as vertical archaeal legacies, are bacterial ‘Serial Innovations’ stabilized in Asgard lineages. Our work extends this trajectory to the translational machinery, traditionally considered the archaeal component of the eukaryotic cell, further demonstrating that the bacterialization of the proto-eukaryote went far beyond a piecemeal accumulation of metabolic or maintenance factors^41^.

While single-gene phylogenies must be interpreted with caution, the CCA-adding enzyme offers a high-resolution lens to trace deep evolutionary events. As a monomeric, stand-alone catalyst, it may be less susceptible to the co-evolutionary constraints that can obscure the signal of multi-subunit complexes. Our identification of an active, BCR-derived CCA2 enzyme alongside an intact but silenced archaeal CCA1 enzyme in Heimdallarchaeaceae (Heim12), marked by a specific catalytic-site substitution, provides evidence for a functional replacement of the ancestral archaeal enzyme.

These results also offer an additional perspective on current topological debates concerning the placement of eukaryotes with regards to Asgard archaea^2,9,10^. Although species-tree proximity remains a subject of intense discussion, with various models favoring Hodarchaeales^9^, or the broader Heimdallarchaeia closest to eukaryotes^2^, our analysis suggests that genomic proximity in a species tree does not always equate to a shared functional ancestry. The divergent, two-copy CCA2 profile of Hodarchaeales may reflect lineage-specific horizontal transfers, or duplications, although the Hodarchaeota CCA2 sequences are more similar to later-diverging bacterial clades which do not contain this eCCA signature found in early branching eukaryotes. In contrast, the stabilized eCCA signature in Heimdallarchaeaceae points toward this lineage as a potentially relevant model for the ancestral cellular framework. This model is further supported by the ecological co-occurrence of Heimdallarchaeaceae and BCR-clade bacteria in bathypelagic marine sediments^40^, providing a plausible physical context for such an ancient transfer.

Ultimately, while the full complexity of eukaryogenesis cannot be resolved by a single gene family, the CCA-adding enzyme phylogeny identifies a reconfiguration of the translational core during a pre-eukaryogenesis phase of innovation. Our results suggest that the eukaryotic ancestor inherited a bacterialized translational system that had already been integrated and stabilized within an archaeal host lineage, one whose affiliation with pre-GOE bacterial signatures places this remodelling event in the anaerobic world, before the emergence of the eukaryotic cell. This has the potential to highlight how functional integration of bacterial-derived innovations in core machinery, rather than just scattered acquisitions, established the functional template for cellular complexity.

## Methods

### Genome Acquisition

**Bacterial genomes** were downloaded from the BV-BRC database (v. 3.53.3)^42^. Representative genomes or high-quality complete assemblies were prioritised to ensure reliable gene annotation. Where possible, three genomes per bacterial lineage were selected, with lineage composition informed by the taxonomic groupings used in Nelson-Sathi et al.⁴² as a reference framework for bacterial diversity relevant to archaeal-bacterial gene flow. Taxonomic assignments were subsequently updated to reflect current GTDB classification (r220)^35^. A total of 80 bacterial genomes were downloaded (see Supplementary Table 3). Additionally, 16 genomes from Patescibacteria (formerly CPR) were obtained from the BV-BRC database to represent ultra-small bacteria with reduced genomes.

**Archaeal genomes** were primarily sourced from the BV-BRC database. To expand sampling of Asgard archaea, particularly Heimdallarchaeia, additional metagenome-assembled genomes (MAGs) reported by Appler et al.^11^, were incorporated into the dataset. Since many archaeal genome sequences derive from MAGs, they exhibit variable quality and are susceptible to contamination and completeness issues. CheckM2 was used to assess genome quality^43^, a tool selected for its accuracy with the reduced genome sizes and unusual genomic features characteristic of archaeal lineages, particularly DPANN and Asgard groups. Only genomes meeting stringent quality thresholds (greater than 85% completeness and less than 5% contamination) were retained for downstream analysis. To identify and remove duplicate assemblies, MD5 hashes were calculated for all genome files, and exact duplicates were excluded (see Supplementary Table 1).

**Early-branching eukaryotic genomes** were downloaded from NCBI by querying each major early branching eukaryotic clade using the taxonomy database, following a recently updated tree of life for major eukaryotic groups^44^. A total of 52 genomes were downloaded, prioritizing reference genomes where available. Partial or low-quality assemblies were excluded. For lineages where genomic sequences were unavailable or failed to yield CCA-adding enzyme homologs, we searched transcriptome shotgun assembly (TSA) databases using tblastn to retrieve transcript sequences of CCA sequences (see Supplementary Table 3).

### CCA-Adding Enzyme Identification

**CCA2 identification in bacteria.** an experimentally validated CCA-adding enzyme from *Bacillus subtilis* strain 168 (UniProt accession P42977)^45^ was used as the query sequence for tblastn searches against all bacterial genomes. This protein has been biochemically characterized and confirmed as a functional Class 2 CCA-adding enzyme. Loci of interest were identified using an E-value threshold of 10⁻⁵, and flanking regions of 300 nucleotides on each side were extracted to ensure capture of complete open reading frames, since the C-terminus is variable and often not captured by blast. Protein sequences were predicted from these genomic regions using Prodigal (v. 2.6.3)^46^.

The predicted protein sequences were aligned using MUSCLE as implemented in AliView (v. 1.0)^47^. We then applied a custom classification pipeline to distinguish genuine CCA-adding enzymes from poly(A) polymerases (PAP) and to identify subfunctionalized variants (CC/A-adding enzymes). This workflow was adapted from established protocols for CCA enzyme classification and relied on the presence or absence of five conserved motifs characteristic of Class 2 CCA-adding enzymes.^26^ This can be found in the GitHub repository linked in the supplement (https://github.com/ellacass/Heimdall_CCA_paper/tree/main). Sequences missing more than one key motif (excluding motif M5, which shows variable conservation) were excluded from further analysis as definitive classification was not possible.

To distinguish CCA-adding enzymes from poly(A) polymerases, sequences were screened for the poly(A) polymerase-specific upstream motif characterized by the amino acid signature [LIV][LIV]G[RK][RK]F.[LIV][AILMVF][HQL][LIV]^19^. Sequences matching this pattern were classified as PAPs and removed. The remaining sequences were further categorized as canonical CCA-adding enzymes (no evidence of subfunctionalization), CC-adding enzymes, or A-adding enzymes based on the presence or absence of flexible loop regions and specific catalytic residues that distinguish these functional variants.^19^ This classification workflow can be seen in Supplementary Figure 6. All bacterial genomes analyzed, including accession numbers, species assignments, processing details, and final CCA enzyme classifications, are provided in Supplementary Table 3.

**CCA2 identification in archaea.** To identify Class 2 CCA-adding enzyme homologs in archaeal genomes, we performed tblastn searches using a *Candidatus* Helarchaeota archaeon protein (NCBI ID: NVM04899.1) as the query. This query protein was initially identified by blasting an *E. coli* protein against the NCBI core nucleotide database, filtered for the archaeal domain (Taxon ID: 2157). Since the presence of CCA2 in archaea had not been previously documented or reported, a permissive E-value threshold of 10⁻⁵ was used. To ensure capture of complete genes, particularly the often-variable C-terminal regions, a flanking region of 500-nucleotide from both sides of each identified locus was extracted for CDS prediction.

All identified loci were then processed through Prodigal^46^ for gene prediction and manually inspected to confirm the presence of full-length open reading frames with sizes and domain structures consistent with CCA-adding enzymes. This two-step approach (permissive initial search followed by stringent validation) was necessary to balance sensitivity and specificity when searching for an enzyme class not previously known to exist in the target genomes. The resulting high-quality CCA2 sequences were used for all downstream phylogenetic analyses.

**CCA2 identification in eukaryotes.** A reference ancestral eukaryotic-type CCA-adding enzyme (eCCA) from *Hartaetosiga balthica* (a choanoflagellate) was used as the query for tblastn searches against early-branching eukaryotic genomes. This approach retrieved CCA-adding enzymes across diverse protistan lineages, several of which contained multiple CCA enzyme copies per genome, a phenomenon previously documented in organisms such as *Acanthamoeba castellanii*, which possesses four distinct CCA-adding enzyme variants.^48^

Flanking regions of 300 nucleotides were extracted for each locus. Complete protein sequences were predicted using Augustus (v 3.5.0) with the Alveolata model^49^, selected for its suitability for protistan gene structures. For sequences where Augustus failed to predict a complete coding sequence, the ExPASy Translate Tool^50^ was used to manually identify open reading frames. To refine exon-intron boundaries and validate sequence accuracy, genomic sequences were aligned against available transcriptome shotgun assemblies (TSA) from NCBI. This manual curation was essential for several lineages with complex gene structures or fragmented assemblies. Sequences requiring manual reconstruction are listed in Supplementary Table 3.

Sequences were inspected for the presence of the five universally conserved Class II catalytic motifs (Motifs A–E). Nine sequences were excluded due to the absence of Motif 4, a critical catalytic element; these were interpreted as potentially non-functional gene fragments or pseudogenes. To avoid redundancy and the overrepresentation of specific taxa, paralogous sequences were pruned. For species with multiple sequence variants (e.g., *Anaeramoeba flamelloides*, *Diacronema lutheri*, and *Entamoeba histolytica*), only the longest variant or the one exhibiting the highest conservation of catalytic residues was retained. Five sequences (e.g., *Isotricha intestinalis* and *Lotmaria passim*) were removed following preliminary phylogenetic analysis due to extreme branch lengths (branch length > 1.0) or unstable, atypical placements that suggested a high risk of Long Branch Attraction (LBA) artifacts. This curation resulted in a final dataset of 24 eukaryotic sequences for analysis.

**CCA1 identification in archaea.** To assess the distribution of Class 1 CCA-adding enzymes across archaea, tblastn searches were performed using the *Archaeoglobus fulgidus* Class 1 enzyme as the query^18^. This protein was selected because its three-dimensional structure has been experimentally determined^18^. Searches were conducted with an E-value threshold of 10⁻¹⁰. Genomic loci were extracted with 500-nucleotide flanking regions, and coding sequences were predicted using Prodigal as described above for CCA2 identification.

### Species Tree Construction

Species trees for archaeal lineages were constructed using REvolutionH-tl (v. 1.4.8)^20^, a graph-based phylogenomic tool that produces reconciled species trees directly from proteomes without relying on predefined marker gene sets. Only archaeal genomes that passed all quality control criteria, meeting contamination and completeness thresholds, lacking exact duplicates, and containing at least one CCA-adding enzyme (CCA1, CCA2, or both), were included in species tree reconstruction. This filtering ensured that the analyzed genomes represented high-confidence archaeal lineages suitable for inferring CCA enzyme evolutionary patterns. The resulting trees were visualized with branches colored according to CCA enzyme composition to reveal the phylogenetic distribution of enzyme types across archaeal superphyla. Supplementary Figure 7 shows pipeline for all filtering steps, gene prediction steps and species tree construction.

### Cloning of archaeal genes

Open reading frames of the archaeal CCA-adding enzymes were codon-optimized for recombinant expression in *E. coli*, elongated by a genomic N- or C-terminal His_6_ tag sequence, and cloned into the pET28a(+) vector by GENEWIZ.

### Recombinant expression and purification of archaeal CCA-adding enzymes

Commercially generated plasmids were used to transform *E. coli* BL21 (DE3) cca:cam cells lacking the endogenous CCA-adding enzyme. Cells were grown at 37 °C in liquid LB medium (supplemented with 50 µg/ml kanamycin and 35 µg/ml chloramphenicol) to an OD_600_ of 1.3 – 1.5. Recombinant expression was induced by adding 0.1 – 1 mM IPTG. After 16 h of incubation at 16 °C, cells were harvested by centrifugation (5,330 rcf for 15 min) and stored at −70 °C until use. Cell pellets were resuspended in ice-cold 25 mM Tris/HCl (pH 7.5), 500 mM NaCl and 1 mM DTT and lysed by homogenization with zirconia beads at 6 m/s for 30 s. The lysate was centrifuged at 30,600 rcf and 4 °C for 30 min and sterile-filtered. After equilibration of a Ni^2+^-loaded HisTrapFF 1ml column (Cytiva) with 5 column volumes of 25 mM Tris/HCl (pH 7.5), 500 mM NaCl and 5 mM imidazole, the lysate was applied to the column. The column was washed with 15 column volumes under increased imidazole concentration (45 mM) followed by a further increase to 500 mM imidazole for enzyme elution with 5 column volumes. Enzyme fractions were dialyzed in ice-cold 25 mM Tris/HCl (pH 7.5) and 200 mM NaCl for 16 h and stored in 10 % (v/v) glycerol at −70 °C. Impure fractions were further purified by size exclusion chromatography on a HiLoad^TM^ Superdex 75 pg column (Cytiva) according to Hennig et al^51^. All enzyme fractions were controlled for purity by SDS-PAGE and stored on 10 % (v/v) glycerol at −70 °C. Protein concentration was determined by measuring UV absorption at 280 nm.

### In vitro transcription of tRNA

tRNA^Phe^ was transcribed *in vitro* in the presence of 0.11 µM radioactively labeled α-^32^P-ATP (3000 Ci/mmol), using a transcription cassette flanked by hammerhead and HDV ribozyme sequences^52^. Transcription was performed in 80 mM Tris/HCl (pH 7.5), 22 mM MgCl_2_, 1 mM spermidine, 8.3 mM DTT, 120 µg/ml acetylated BSA and 4 mM NTPs with 0.4 U thermostable inorganic pyrophosphatase (TIPP, NEB) and recombinant T7 RNA polymerase (40 ng/µl). Ribozyme activity ensured generation of homogenous 5’-OH- and 2’-3’-tRNA ends. The resulting tRNAs were purified by denaturing 10% PAGE and eluted in 200 mM NaCl, 1 mM EDTA and Tris/HCl (pH 7.5) at 65 °C, followed by ethanol precipitation. tRNAs were 5′-phosphorylated and 3′-dephosphorylated before use.

### Enzyme activity tests

100 - 200 ng of purified enzymes were incubated with 5 pmol of radioactively labeled tRNA^Phe^, 50 mM glycine/NaOH (pH 8.5), 10 mM MgCl_2_, 2 mM EDTA, 1 mM NTPs, 2 mM DTT, 4 mM spermidine and 7.5 pmol acetylated BSA for 5 - 30 min at 37 °C. After ethanol precipitation, samples were separated by denaturing PAGE and analyzed by autoradiography.

### BLASTp analysis and heatmap visualization

To assess large-scale similarity relationships among CCA2 proteins independently of phylogenetic reconstruction, pairwise BLASTp (v. 2.16.0)^53^ analyses were performed with an E-value cutoff of 10⁻¹⁰ using full-length protein sequences. Two complementary analyses were conducted. In the first, Heimdallarchaeia CCA2 sequences (Heim12 and Heim02) were queried against a reference dataset consisting of bacterial and early-branching eukaryotic CCA2 proteins. In the second, early-branching eukaryotic sequences were incorporated into the query set alongside Heimdallarchaeia sequences and queried against the same bacterial reference dataset. Since CCA2 proteins exhibit substantial variation in total length due to divergent N- and C-terminal regions, similarity scores were normalized to reduce length-associated bias. For each BLASTp hit, a normalized similarity score (*S*) was calculated as:

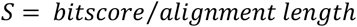

where alignment length corresponds to the homologous region aligned by BLASTp. Since BLASTp does not force-align non-homologous terminal extensions, this normalization captures conservation density within shared aligned regions rather than total protein size. To independently assess alignment completeness, coverage (*C*) was calculated as:

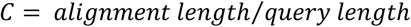

Coverage values were used exclusively for quality annotation and visualization. Alignments with high coverage but disproportionately weak similarity scores (*W*) were classified as probable extended low-information matches using the following criteria:

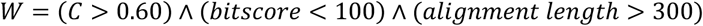

These alignments typically correspond to broad but weakly conserved matches in which BLASTp extends alignments across poorly conserved regions and can be seen in Fig. 7 as having grey cells.

Because normalization by alignment length can artificially inflate normalized similarity values for short alignments with moderate bitscores, an additional annotation category was introduced to identify conserved partial matches (*P*). These were defined as:

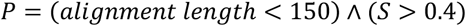

These hits that match these criteria can be seen in Fig. 7 as red cells.

For each query–subject pair, the maximum normalized similarity score was retained and converted into a similarity matrix. Hierarchical clustering was performed independently on rows and columns using Spearman correlation-based distance (1 - ρ) and complete-linkage agglomerative clustering implemented in the pheatmap package in R (v. 2025.09.0+387)^54^. Clustering therefore reflects similarity in overall BLAST profile structure across the dataset rather than direct phylogenetic inference.

### Bacterial gene tree construction

Phylogenetic reconstruction was performed using IQ-TREE 3 (v3.0.1)^37^. The analysis was based on an amino acid alignment of 104 sequences containing 846 sites, of which 569 were parsimony-informative, 169 were singletons, and 107 were constant. The optimal evolutionary model was selected via ModelFinder using the Bayesian Information Criterion (BIC). The Q.PFAM+I+R6 model was identified as the best-fit model, accounting for a proportion of invariable sites (+I) and site-specific rate heterogeneity using a FreeRate model with six rate categories (+R6). Maximum likelihood tree search was initiated from a random seed (524442). Branch support was evaluated using 1,000 ultrafast bootstrap replicates (UFBoot) and 1,000 replicates of the SH-like approximate likelihood ratio test (SH-aLRT), applied simultaneously. The tree was rooted using Aquificota as outgroup, consistent with its established position as one of the deepest-branching bacterial lineages. This rooting strategy was further supported by two independent rooting approaches: midpoint rooting and minimal ancestor deviation (MAD)^55^ rooting both independently placed the root within the CC-adding enzyme clade, consistent with the outgroup-defined topology.

### All taxa gene tree construction

To investigate the broader evolutionary history of the CCA-adding enzyme across domains, the same input as used for the heatmap was used. To reduce sequence redundancy while maintaining phylogenetic diversity, sequences were clustered using CD-HIT, with a sequence identity threshold of 95%. After clustering and quality filtering, the dataset consisted of 164 sequences. Alignments were processed using the KPIC-Gappy flag in ClipKit^56^. This filtering approach was used to maximise phylogenetic signal and minimise noise. We retained only Parsimoniously Informative and Constant (KPIC) sites, defined as columns containing at least two amino acids across at least two taxa, while also retaining conserved positions. To address alignment sparsity, a Gappy threshold of 0.5 was applied, excluding any site where more than 50% of the taxa contained gaps. This bioinformatic trimming was supplemented by manual removal of the non-conserved N-terminal (∼155 sites) and C-terminal (∼545 sites), restricting the analysis to the catalytic core to mitigate compositional bias and saturation. A breakdown of the extent to which compositional bias was reduced at each threshold can be found in Supplement Data 1.

### Maximum Likelihood

The maximum-likelihood gene tree was constructed using IQ-TREE3 (v3.0.1)^37^. To mitigate the impact of alignment uncertainty and compositional bias on phylogenetic inference, we systematically evaluated various gap thresholds using KPIC-Gappy (from 0.9 to 0.5). At less stringent thresholds (0.9 to 0.6), the optimal substitution model selected by ModelFinder remained anchored to the VT empirical matrix, and a high proportion of sequences (>61%) continued to fail the chi-squared test for compositional homogeneity. Applying a 0.5 threshold triggered a shift in the optimal substitution model from VT to Q.PFAM+I+R6. Because the Q.PFAM matrix is specifically estimated from conserved protein domains, this shift indicates that the 0.5 threshold successfully removed noisy, rapidly evolving, and compositionally biased sites, revealing the conserved phylogenetic core of the protein. Combining this threshold with manual tail removal reduced the proportion of sequences failing the compositional homogeneity test to 34.4%. This optimally trimmed alignment was therefore selected for final tree reconstruction. To further account for across-site compositional heterogeneity and mitigate long-branch attraction artefacts inherent to deep-time phylogenomics, we applied the Posterior Mean Site Frequency (PMSF) approximation to the mixture model. The maximum likelihood topology inferred under Q.PFAM+I+R6 served as the guide tree for PMSF site-frequency estimation. Site-specific amino acid frequency profiles were calculated from this guide tree and fixed for a full maximum likelihood tree search, correcting for mutational saturation at rapidly evolving positions. Branch support was evaluated on the optimised topology using 1,000 ultrafast bootstrap replicates (UFBoot) and 1,000 replicates of the SH-like approximate likelihood ratio test (SH-aLRT), applied simultaneously.

To evaluate whether alternative arrangements of the three focal groups could be statistically rejected, topology testing was performed using IQ-TREE 3. Full trees representing all three possible unrooted topologies for the BCR, Heim12, and eukaryote clades: (i) BCR sister to (Heim12 + eukaryotes), (ii) Heim12 sister to (BCR + eukaryotes), and (iii) eukaryotes sister to (Heim12 + BCR) were generated by rearranging the PMSF bootstrap tree while preserving all other taxon relationships and branch lengths. Log-likelihoods were evaluated under the Q.PFAM+I+R6 model using the approximately unbiased (AU) test (-z −au −zb 10000 −n 0).

### Bayesian methods

Bayesian phylogenetic inference was performed using MrBayes (v3.2.7)^57^. To account for uncertainty in the amino acid substitution process without *a priori* model selection, we employed a mixed-model strategy (prset aamodelpr=mixed), allowing the MCMC to sample across fixed-rate matrices. Rate variation among sites was modeled using a gamma distribution with four categories. Initial analyses using two independent runs of four chains each (one cold, three heated) showed poor convergence. To improve exploration of tree space, we extended the analysis to 10,000,000 generations and increased to eight chains per run with a reduced temperature parameter. Despite this, the two independent runs failed to converge: run 2 underwent a single large transition of approximately −8,000 log-likelihood units, reaching a final log-likelihood of approximately −38,000, while run 1 was unable to locate this region of tree space. The resulting ASDSF remained high, reflecting a genuinely multimodal posterior landscape for this short, deeply divergent alignment rather than insufficient sampling effort. Since the runs did not converge, posterior probabilities from a combined consensus are not reported as quantitative support. Post-burn-in trees were extracted from run 2 only (2,500 trees discarded as burn-in), as this run demonstrably occupied the higher-likelihood region, and its topology was used solely for qualitative comparison with maximum-likelihood reconstructions. Phylogenetic conclusions are therefore based on concordance among ML analyses, site-filtered alignments, with the run 2 Bayesian topology serving as independent corroboration.

## Data availability

The data supporting the findings of this study are available within the paper and its Supplementary Information files, or from public databases as detailed below:

All bacterial and archaeal genomic sequences analyzed in this study were retrieved from the BV-BRC database (v. 3.53.3). Additional Archaeal genome assemblies used for comparative analysis were obtained from Appler et al. (2026)^11^ and are accessible via the original publication’s supplementary data. All genomes used in final datasets are outlined in Supplementary Table 1.

Primary query sequences for homolog identification are publicly available via the UniProt Knowledgebase and NCBI Protein database under accession IDs: P42977 (*Bacillus subtilis* strain 168), NVM04899.1 (*Candidatus* Helarchaeota archaeon).

## Code availability

Reconciled species trees and proteome-wide orthogroup inferences were generated using the graph-based phylogenomic tool REvolutionH-tl (v.1.4.8), available at https://pypi.org/project/revolutionhtl/

Open reading frame predictions were conducted via Prodigal (v.2.6.3), alignment trimming utilized ClipKit, model selection was handled by ModelFinder, and maximum-likelihood trees were executed using IQ-TREE 3.

Downstream data parsing and filtering scripts used for sequence analysis have been archived on GitHub at [https://github.com/ellacass/Heimdall_CCA_paper/tree/main].

## Acknowledgements

We thank Stephan Bernhart for proofreading the manuscript, and also Markus Lechner for initial bioinformatical analyses of archaeal class 2 coding sequences.

## Funding

This work was supported by the Deutsche Forschungsgemeinschaft (DFG, German Research Foundation) through project grant [PR 1288/10-1] to S.J.P., and project grant [MO 634/25-1] to M.M.

E.C. is supported by the School for Embedded and Composite Artificial Intelligence (SECAI), funded by the German Federal Ministry of Education and Research (BMBF) and the German Academic Exchange Service (DAAD).

## Author contributions

Ella Cassidy and Claudius Doktor contributed equally to this work. Correspondence and requests for materials should be addressed to Ella Cassidy.

S.J.P., M.M and H.B conceived and designed the study. E.C. implemented the computational framework, curated the genomic datasets, and performed all phylogenomic and phylogenetic analyses. C.D., H.B., and M.M. designed, performed, and supervised the biochemical assays and functional experiments. E.C., C.D., H.B., M.M., and S.J.P. analyzed the structural and evolutionary data. E.C. wrote the initial manuscript draft. All authors contributed to data interpretation, reviewed the manuscript, and approved the final text.

## Competing interests

There are no competing interests to declare.

## Notes

https://github.com/ellacass/Heimdall_CCA_paper/tree/main

### Competing Interest Statement

The authors have declared no competing interest.

### Summary of Updates

Title revised; Extra supplementary figure; figures updated

