## Supplementary Figures 1-7 for "Found in translation: CCA-adding enzymes reveal bacterialization of Asgard archaea prior to eukaryogenesis"

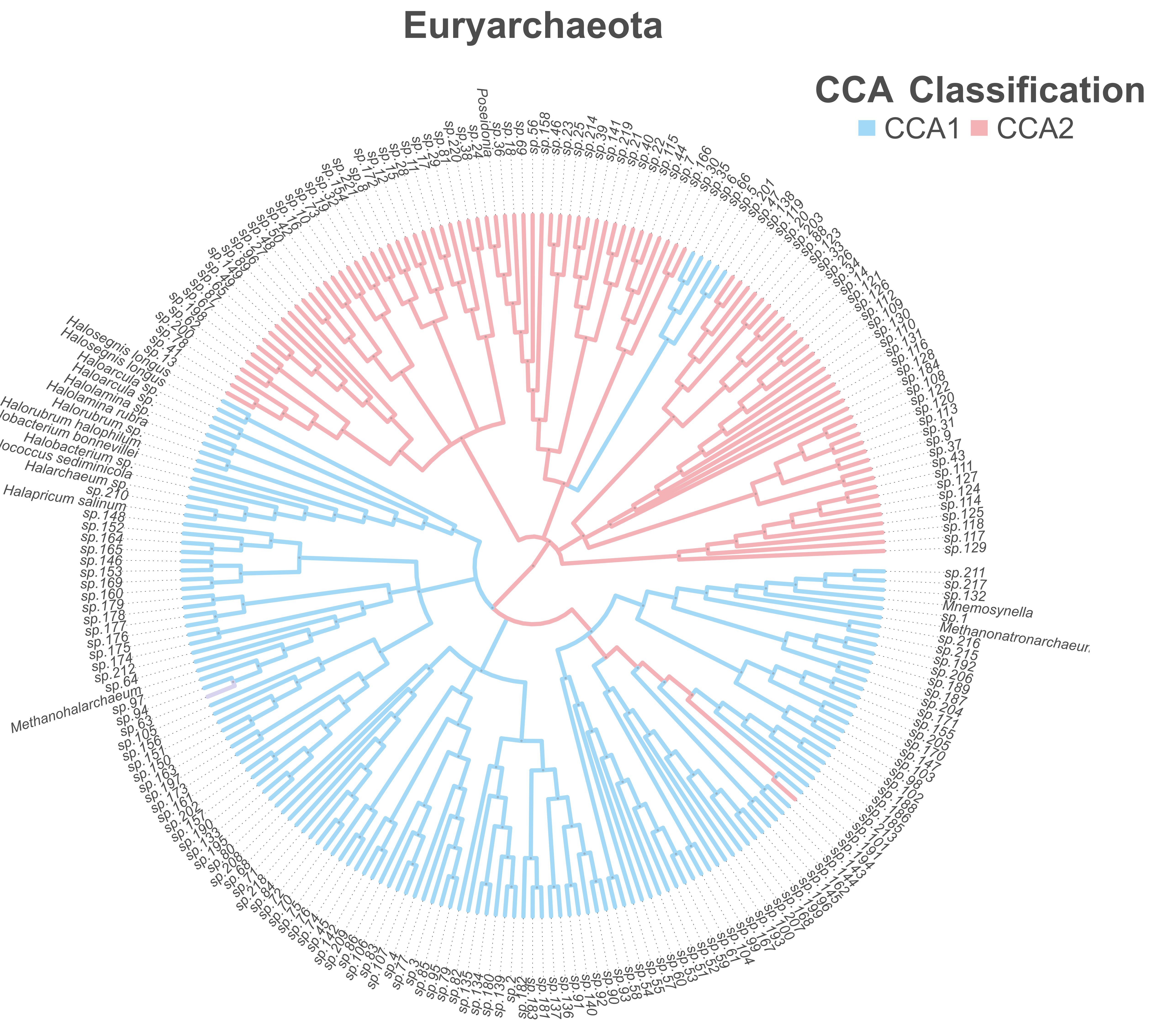


**Supplementary Figure 1**. Species tree of Euryarchaeota constructed using REvolutionH-tl from proteomes. Branches are coloured by CCA-adding enzyme class; blue genomes contain archaeal CCA1, whereas pink genomes contain bacterial-type CCA2. Accessions relating to sp labels can be found in Supplemental table 1.

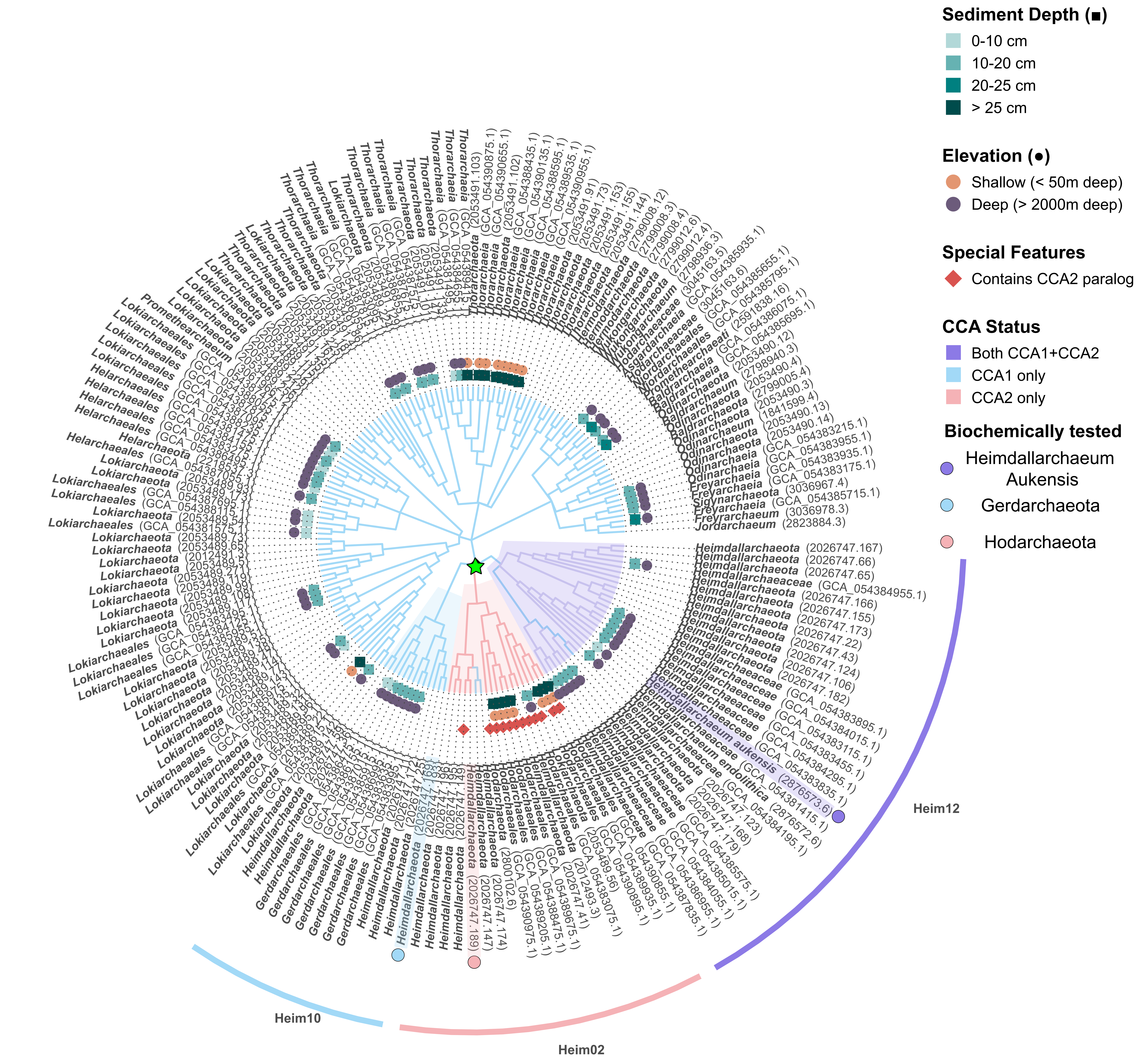


**Supplementary Figure 2.** Species tree of Asgard archaea. Supplementary Figure 2. Species tree of the Asgard superphylum reconstructed with REvolutionH-tl, as in Figure 4, with genome accession numbers included at leaf labels. Branches are coloured by CCA-adding enzyme class: blue, CCA1 only; red, CCA2 only; purple, both CCA1 and CCA2. Metadata from Appler et al.

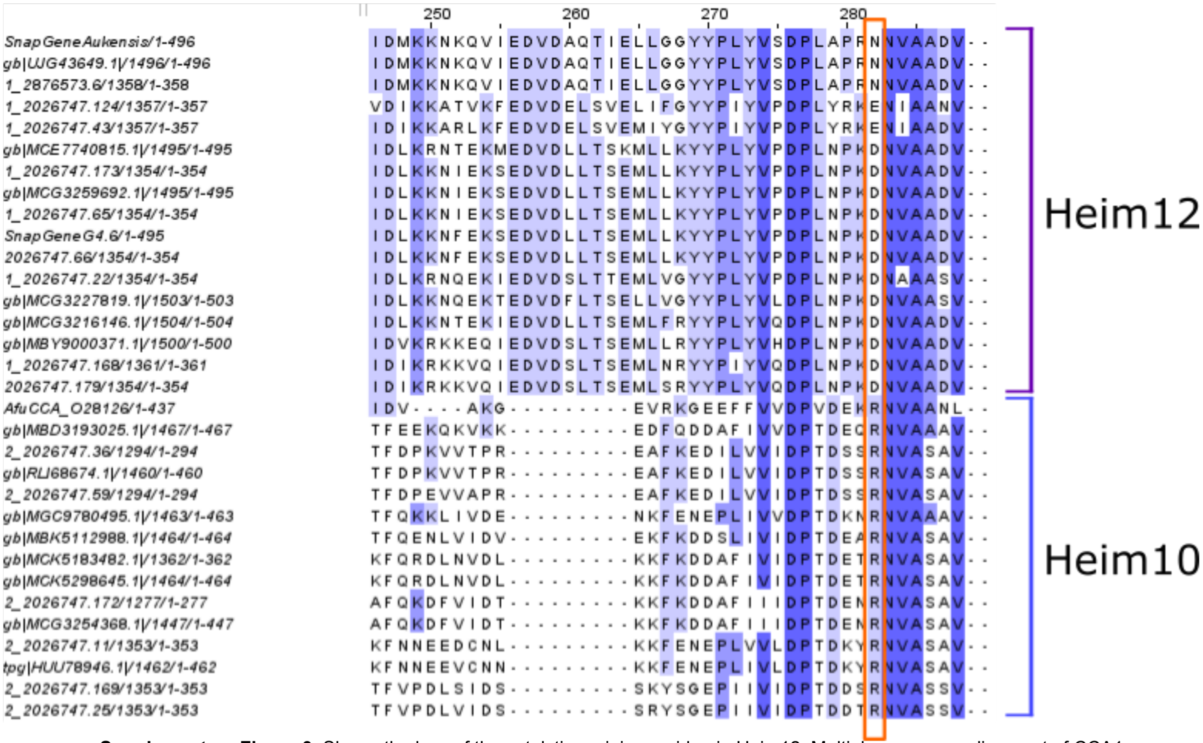


**Supplementary Figure 3**. Shows the loss of the catalytic arginine residue in Heim12. Multiple sequence alignment of CCA1 sequences from Heim12 (Heimdallarchaeaceae) and Heim10 (Gerdarchaeales) across alignment positions 250–285. In canonical class 1 CCA-adding enzymes, a highly conserved arginine residue plays a critical role in CTP and ATP substrate selection during CCA end synthesis. This arginine is retained across all Heim10 CCA1 sequences, but is consistently absent in Heim12 CCA1 sequences (orange box), where it is replaced by asparagine or a functionally distinct residue. Loss of this residue across the entire Heim12 lineage is consistent with inactivation of the class 1 enzyme, supporting the inference that CCA2 has functionally replaced the ancestral archaeal enzyme in this clade

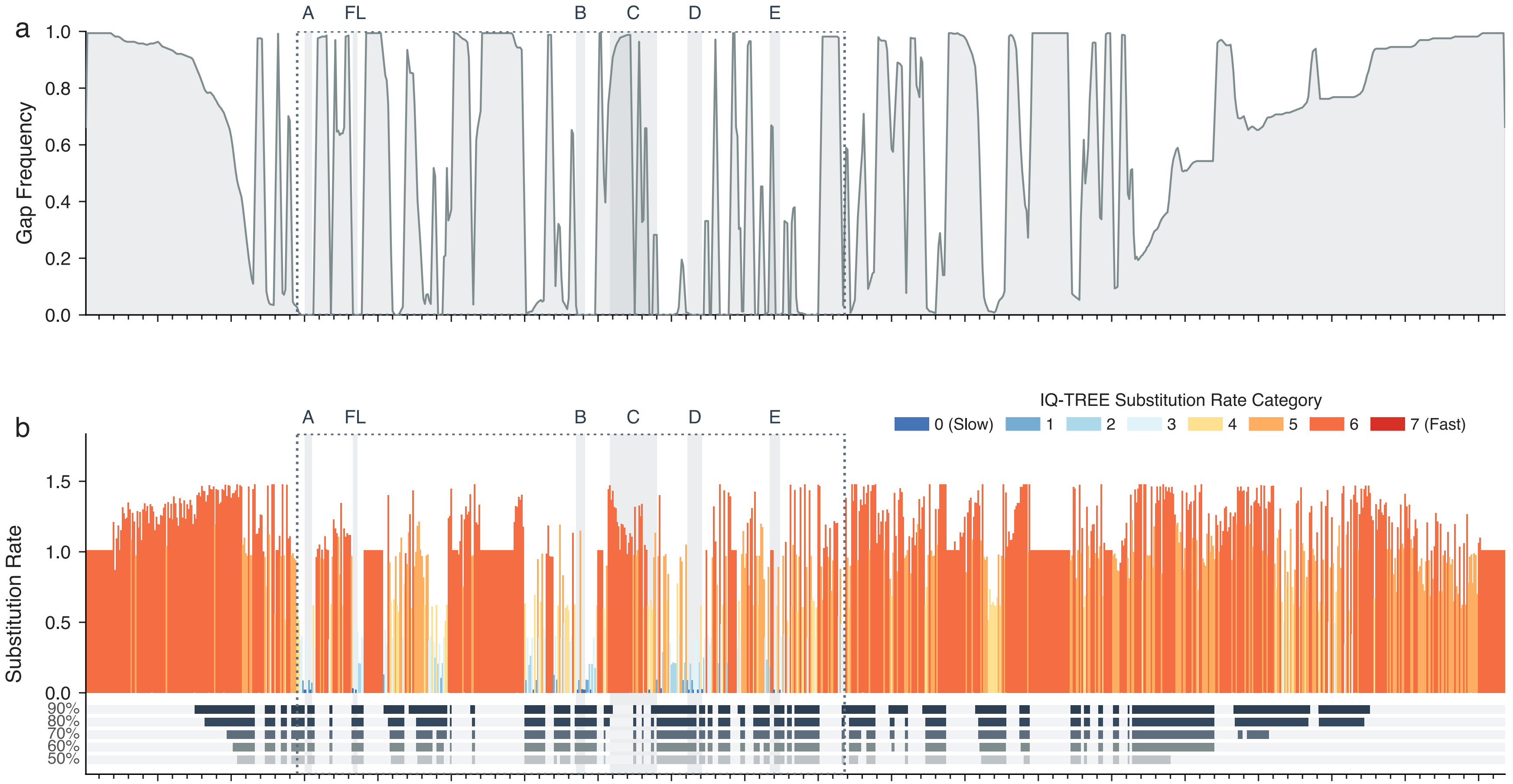


**Supplementary Figure 4.** Site-specific evolutionary profiles, alignment trimming dynamics, and phylogenomic signal mapping for the CCA2 enzyme**.** The dotted bounding box shows the catalytic core (alignment positions 145–518), housing functional motifs A to E and the flexible loop (FL). (a) Multiple sequence alignment (MSA) gap occupancy profile across the full-length enzyme, highlighting near-complete gap saturation across the flanking terminal loops. (b) Site-specific amino acid substitution rates inferred under the IQ-TREE, color-coded by a rate as decided by ModelFinder. Slow-evolving sites (Category 0; blue) to variable, saturated positions (Category 7; crimson). The horizontal alignment tracks below demonstrate the strict retention profiles across progressive ClipKIT kpic-gappy stringency thresholds (90% down to 50%).

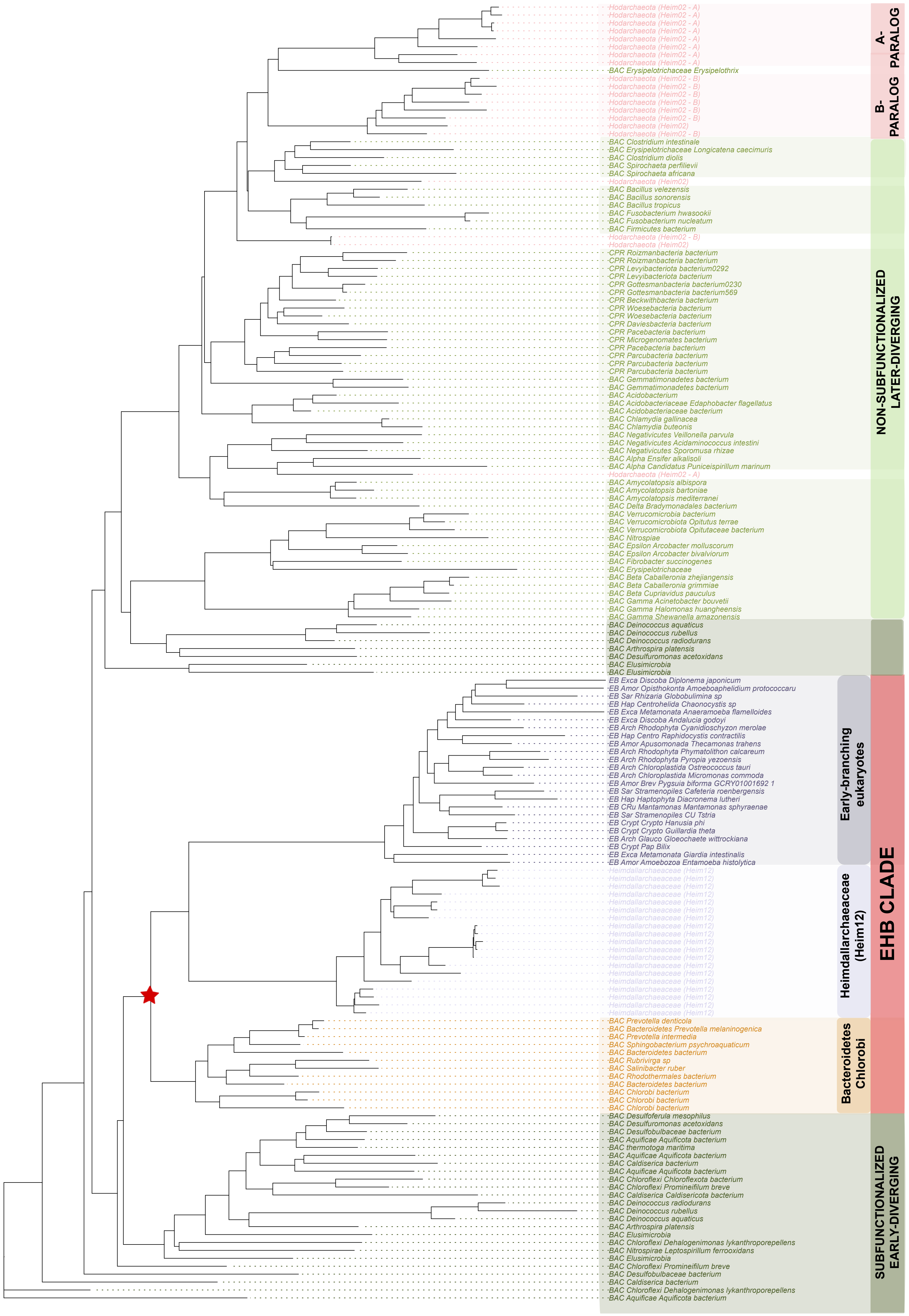


**Supplementary Figure 5**. Bayesian inference was performed using MrBayes (v3.2.7) with a mixed amino acid substitution model (prset aamodelpr=mixed) and gamma-distributed rate variation across four categories. Two independent runs of eight chains each were conducted for 10,000,000 generations; due to non-convergence between runs (elevated ASDSF; run 2 underwent a single large likelihood transition of approximately −8,000 log-likelihood units), posterior probabilities are not reported as quantitative support. The topology shown is derived from post-burn-in trees of run 2 only (2,500 trees discarded as burn-in), which occupied the higher-likelihood region (final log-likelihood ≈ −38,000), and is presented solely for qualitative comparison with the maximum-likelihood reconstruction (Fig. 8 in the main text). The monophyly of the EHB clade (early-branching eukaryotes, Heimdallarchaeaceae, and Bacteroidetes/Chlorobi/Rhodothermota) was independently recovered in this Bayesian topology, consistent with the maximum-likelihood result.

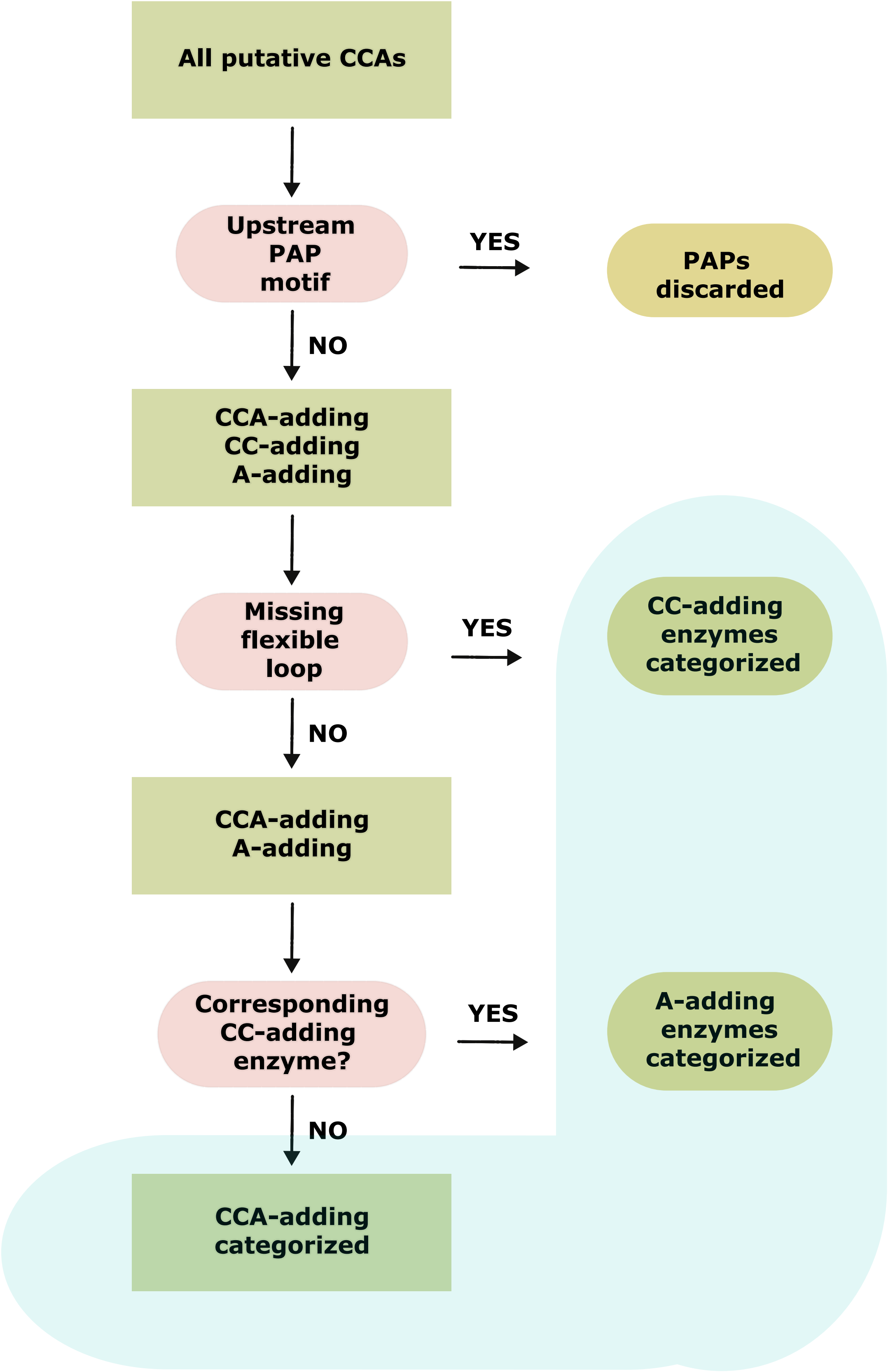


**Supplementary Figure 6**. Classification workflow for distinguishing CCA-adding, CC-adding, A-adding, and poly(A) polymerase (PAP) enzymes. All putative bacterial Class 2 CCA-adding enzyme sequences identified by tblastn were passed through a sequential decision pipeline. First, sequences were screened for the upstream PAP-specific motif ([LIV][LIV]G[RK][RK]F.[LIV][AILMVF][HQL][LIV]); sequences matching this signature were classified as poly(A) polymerases and discarded. Remaining sequences were assessed for the presence of the flexible loop element: sequences lacking this structural feature were classified as CC-adding enzymes (teal). Sequences retaining the flexible loop were then examined for the presence of a corresponding A-adding enzyme in the same genome. Sequences retaining the flexible loop with no corresponding A-adding enzyme in the genome were classified as canonical CCA-adding enzymes (teal). This workflow was applied to all bacterial and archaeal sequences prior to phylogenetic analysis and is described in full in the Methods.

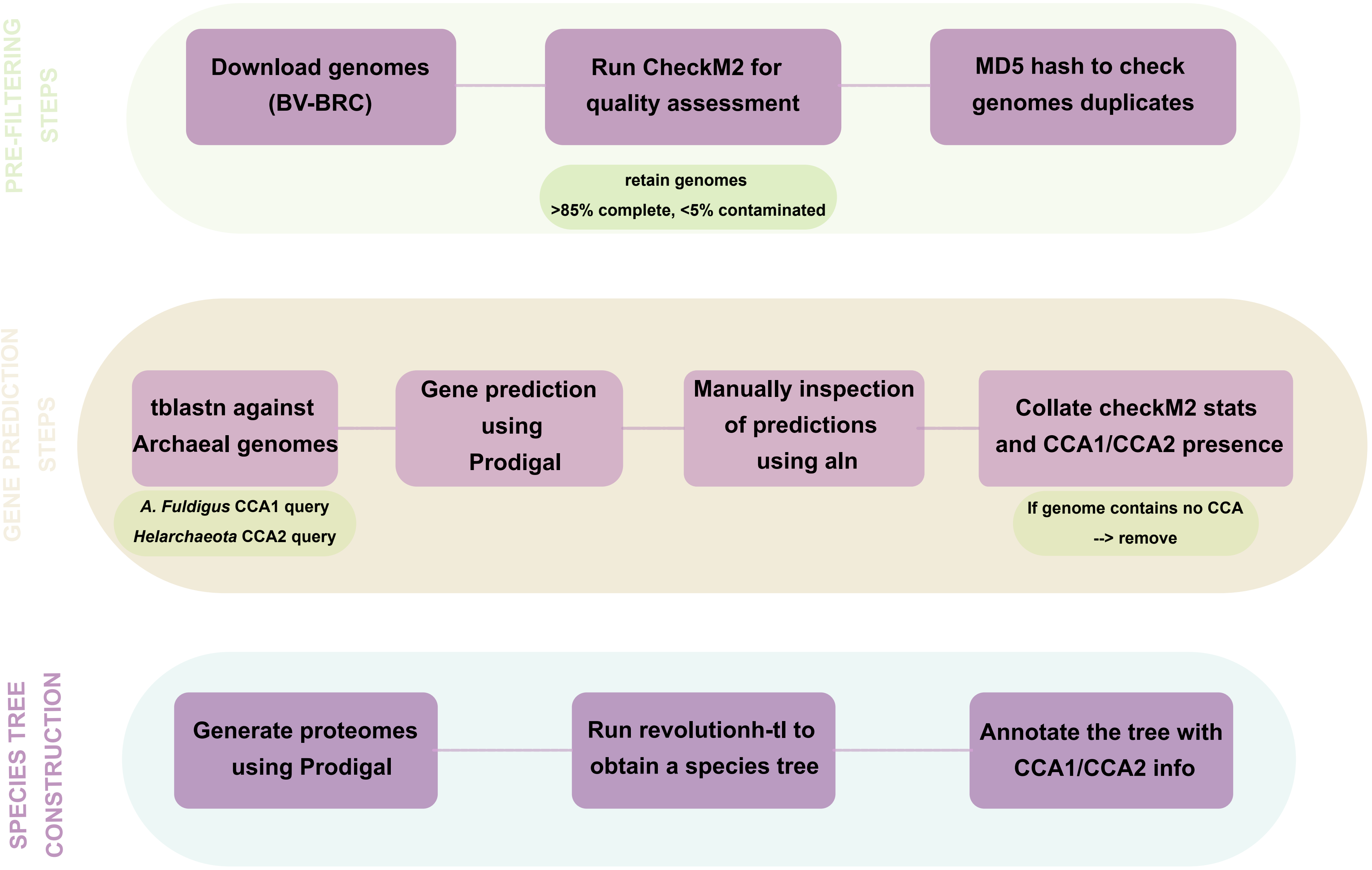


**Supplementary Figure 7. Overview of the archaeal genome processing and species tree construction pipeline.** The workflow is divided into three sequential stages. Pre-filtering steps (top): archaeal genomes were downloaded from the BV-BRC database and assessed for quality using CheckM2; only genomes meeting minimum thresholds of >85% completeness and <5% contamination were retained. Duplicate assemblies were identified and removed using MD5 hash comparison. Gene prediction steps (middle): retained genomes were searched for CCA-adding enzyme homologs using tblastn, with the Archaeoglobus fulgidus CCA1 enzyme and a Candidatus Helarchaeota CCA2 enzyme as queries. Candidate loci were processed through Prodigal for gene prediction and manually inspected using alignment visualisation. CheckM2 quality statistics and CCA1/CCA2 presence calls were collated; genomes containing no detectable CCA-adding enzyme of either class were excluded from downstream analysis. Species tree construction (bottom): proteomes for all retained genomes were generated using Prodigal and used as input for REvolutionH-tl to reconstruct reconciled species trees. Resulting trees were annotated with CCA1/CCA2 enzyme presence information to visualise phylogenetic distribution across archaeal lineages.
